# Novel function of *Hox13* in regulating outgrowth of the newt hindlimb bud through interaction with *Fgf10* and *Tbx4*

**DOI:** 10.1101/2024.11.12.623306

**Authors:** Sayo Tozawa, Haruka Matsubara, Fumina Minamitani, Yasuhiro Kamei, Misako Saida, Momoko Asao, Ken-ichi T. Suzuki, Masatoshi Matsunami, Shuji Shigenobu, Toshinori Hayashi, Gembu Abe, Takashi Takeuchi

## Abstract

5’*Hox* genes regulate pattern formation along the axes of the limb. Previously, we showed that *Hoxa13/Hoxd13* double mutant newts lacked all digits of the forelimbs during development and regeneration, showing that newt *Hox13* is necessary for digit formation in development and regeneration. In addition, we found another unique phenotype. A part of the *Hox13* crispant newts showed hindlimb defects, in which whole or almost whole hindlimbs were lost, suggesting a novel function of *Hox13* in limb development. Using germline mutants, we showed that mutation in *Hox13* led to hindlimb defects. The limb buds of *Hox13* crispants formed, however, did not show outgrowth. Expression of *Fgf10* and *Tbx4*, which are involved in limb outgrowth, decreased in the hindlimb buds of *Hox13* crispants. In addition, hindlimb defects were observed in both *Fgf10* and *Tbx4* crispant newts. Finally, *Fgf10* and *Tbx4* interacted with *Hox13* genetically. Our results revealed a novel function of *Hox13* in regulating the outgrowth of the newt hindlimb bud through interaction with *Fgf10* and *Tbx4*.

## 1 INTRODUCTION

*5’Hox* genes regulate pattern formation along the limb axes. Previous studies using mouse genetics have suggested that the patterns of the stylopod, zeugopod, and autopod regions in the limb are specified by the *5’ Hox* genes, *Hox9/10*, *Hox11*, and *Hox13*, respectively (Fromental-Ramain, Warot, Lakkaraju, et al., 1996; Fromental-Ramain, Warot, Messadecq, et al., 1996; Wellik & Capecchi, 2003; Xu & Wellik, 2011).

In the case of *Hox13*, *Hoxa13^-/-^*, and *Hoxd13^-/-^*, double knock-out mice lost all digits of both the forelimbs and hindlimbs (Fromental-Ramain, Warot, Messadecq, et al., 1996). To determine whether any differences were present in the functions of other tetrapods and to explore the functions during limb regeneration, we generated and analyzed *Hox13* mutant newts. *Hoxa13/Hoxd13* double mutant newts lacked all digits of the forelimbs during development and regeneration, showing that newt *Hox13* is necessary for digit formation during development and regeneration (Takeuchi et al., 2022).

In addition to the loss of whole digits, we found another unique phenotype. A part of the *Hox13* mutant newts showed hindlimb defects (HLDs), in which whole or almost whole hindlimbs were lost. No studies have reported HLDs in *Hox13* knock-out animals. The involvement of *Hox13* in the development of the whole hindlimb is a new and important topic, because *Hox13* has been assumed to work in the autopod of the limb.

The process of early limb development comprises three steps: specification of the limb field, limb bud formation within the field, and outgrowth of the limb bud, with some steps potentially overlapping. Initially, a hindlimb-forming field is specified on the lateral plate mesoderm (LPM) at the cloacal level. The specification of the hindlimb field is likely to be dependent on the *Hox* gene expression; however, unlike the forelimb, no direct evidence is available for the hindlimb (Moreau et al., 2019). The specified LPM cells then form a protruding structure known as a limb bud. Subsequently, the limb bud cells continue to proliferate, leading to outgrowth (Capdevila & Izpisúa Belmonte, 2001). Outgrowth of hindlimb buds is assumed to be regulated by several genes, including *Fgf10, Tbx4,* and *Pitx*1*. Fgf10* signaling is one of the key factors for limb field specification or limb bud formation within the LPM. After *Fgf10* expression starts, *Fgf10* induces *Fgf8* expression in the apical ectoderm and forms the Fgf10-Fgf8 feedback loop. *Tbx4* regulates hindlimb outgrowth by activating *Fgf10* (Naiche & Papaioannou, 2003, 2007). *Pitx1* determines hindlimb identity and regulates *Tbx4* expression in mice (Logan & Tabin, 1999).

In the present study, we observed that the limb buds of *Hox13* crispants formed, however, did not show outgrowth, indicating that *Hox13* regulates the outgrowth of hindlimb buds. We also found that the expression of *Fgf10* and *Tbx4* decreased in the limb buds of *Hox13* crispants. In addition, HLDs were observed in either *Fgf10* or *Tbx4* crispants. Finally, *Fgf10* and *Tbx4* interacted with *Hox13* genetically. These results showed a novel function of *Hox13* in regulating the outgrowth of hindlimb buds by interacting with *Fgf10* and *Tbx4*.

## 2 MATERIALAND METHODS

### 2.1 Newts

In the present study, we used Iberian ribbed newts (*Pleurodeles waltl)* that were raised in our laboratory. The animals were reared as previously described (Hayashi et al., 2013). All procedures were performed in accordance with the Institutional Animal Care and Use Committee of Tottori University (Tottori, Japan) and the national guidelines of the Ministry of Education, Culture, Sports, Science and Technology of Japan.

### 2.2 Preparation of RNPs and microinjection

gRNAs were designed using CRISPRdirect (Naito, Hino, Bono, & Ui-Tei, 2014). The positions of the targets and the sequences are shown in Takeuchi et al., 2022; Suzuki et al., 2024; and Tables S1 and S6. *Fgf10* #3 and # 1 gRNAs in Suzuki et al., 2024 were used as the #1 and #2 gRNAs in this study, respectively. Comprehensive transcriptome data for *P. waltl* showed that no gRNAs had high degrees of identity with other gene bodies (Matsunami et al., 2019). The synthetic tracrRNA, gRNA, and Cas9 protein were obtained from Integrated DNA Technologies (IDT, Coralville, USA). The tracrRNA and gRNA were annealed and Cas9 RNPs were produced following the manufacturer’s instructions, immediately before injection. Microinjection of RNPs was performed as previously described (Hayashi et al., 2019; Suzuki et al., 2018). Additionally, we used FemtoJet (Eppendorf Japan, Tokyo, Japan) for microinjection.

### 2.3 Germline mutants

For generating *Hox13* germline mutants, we first outcrossed *a13/c13/d13* crispants with wild-type newts and obtained three types of triple monoallelic F1 newts (A: *Hoxa13^+/Δ6^*, *Hoxc13^+/Δ8^* and *Hoxd13^+/Δ84^*; B: *Hoxa13^+/Δ40^*, *Hoxc13^+/Δ6^*, and *Hoxd13^+/Δ9.1k^*; and C: *Hoxa13^+/Δ40^*, *Hoxc13^+/Δ6^*, and *Hoxd13^+/Δ1.4k^*). In addition, two types of F2 newts (D: *Hoxa13 ^Δ40/Δ40^*, *Hoxc13 ^Δ6/Δ6^*, and *Hoxd13 ^+/Δ9.1k^*; and E: *Hoxa13^+/Δ40^*, *Hoxc13 ^Δ6/Δ6^*, and *Hoxd13 ^Δ1.4k/Δ9.1k^*) were obtained by crossing F1 type B and type C newts. These two types of F1 and three types of F2 newts were used as parents (Tables 1 and S2). The nucleotide and amino acid sequences of *Hoxa13^Δ40^* and *Hoxc13 ^Δ6^* were reported previously (Takeuchi et al., 2022). The sequences or predicted deletion regions of other mutant alleles are indicated in Figure S1. We used previously reported *Fgf10* germline mutants (Suzuki et al., 2024).

### 2.4 Genotyping using next-generation sequencing

Genotyping was performed as described previously (Takeuchi et al., 2022), except that the Takara Ex Taq Hot Start Version (Takara Bio, Kusatsu, Japan) was used. The target regions were amplified with the primer sets shown in Table S3. Sequencing data were analyzed as previously described (Iida et al., 2020; Suzuki et al., 2018).

### 2.5 Micro-computed X-ray tomography

The 3D raw data were acquired using an R_mCT2 (Rigaku, Tokyo, Japan). The X-ray tube was set at 90 kV and 160 µA. The 3D images were visualized using the 3D Visualization Software OsiriX (Pixmeo SARL, Geneva, Switzerland).

### 2.6 Bone staining

All larval newts were fixed overnight in 70% ethanol at 4°C. The bone and cartilage were stained with Alizarin Red and Alcian Blue according to the following protocol. First, the viscera were removed with a knife and forceps, and gradually hydrated. The samples were bleached with 6% H_2_O_2_ in light conditions overnight at 4°C, which was gradually substituted with 100% ethanol. Subsequently, the samples were rinsed with 70% ethanol and 30% acetic acid. Cartilages were stained using 0.01% Alcian Blue (A5268, Sigma-Aldrich) for 3 to 7 days. The samples were gradually hydrated and rinsed with 0.1 M KOH for 30 min. The bones were then stained with 0.01% Alizarin Red for 1 h, rinsed with 0.1 M KOH for 30 min, and rinsed with DW until the liquid became colorless. Finally, the liquid was substituted by 100% glycerol gradually.

### 2.7 Whole-mount in situ hybridization

Newt embryos were fixed and processed for whole-mount in situ hybridization (WISH) as described previously (Takeuchi et al., 2022). cDNA fragments for antisense and sense probes were obtained from *P. waltl* cDNA (st27) by reverse transcription PCR. Primers are listed in Table S3. Samples were hybridized simultaneously for each gene and the staining reactions were terminated simultaneously. Images were similarly adjusted for each gene.

### 2.8 Statistical analysis

Experimental data were analyzed using Welch’s two-sample-test and Fisher’s exact tests. Values of P<0.05 were considered statistically significant.

## 3 RESULTS

### 3.1 Hindlimb defects in *Hox13* crispants

Newts have four *Hox13* paralogs (*a13, b13, c13*, and *d13*). We previously generated *Hox13* mutants (hereafter referred to as *Hox13* crispants) with high mutation rates (mean of 14 target sites, 97.4%) using CRISPR-Cas9 (Takeuchi et al., 2022). The *Hoxa13/Hoxd13* double mutant newts lacked all digits of the forelimbs (Takeuchi et al., 2022) and hindlimbs (for examples, see Figure 1(a) and 1(c)) during development. We found that a part of newts in both *Hox a13/d13* double and *Hoxa13/c13/d13* triple crispants had hindlimb defects (hereafter referred to as HLDs). Most mutant newts with HLDs lost their whole hindlimbs (hereafter referred to as complete HLD) (for example, see Figure 1(a), white arrowheads). Some mutant newts with HLDs had only small swellings (hereafter referred to as severe HLD) (for example, see Figure 1(a), yellow arrowheads). Mutant newts with hindlimb regions other than the digits were defined as those without HLDs (for example, see Figure 1(c), Without HLDs).

**FIGURE 1.**
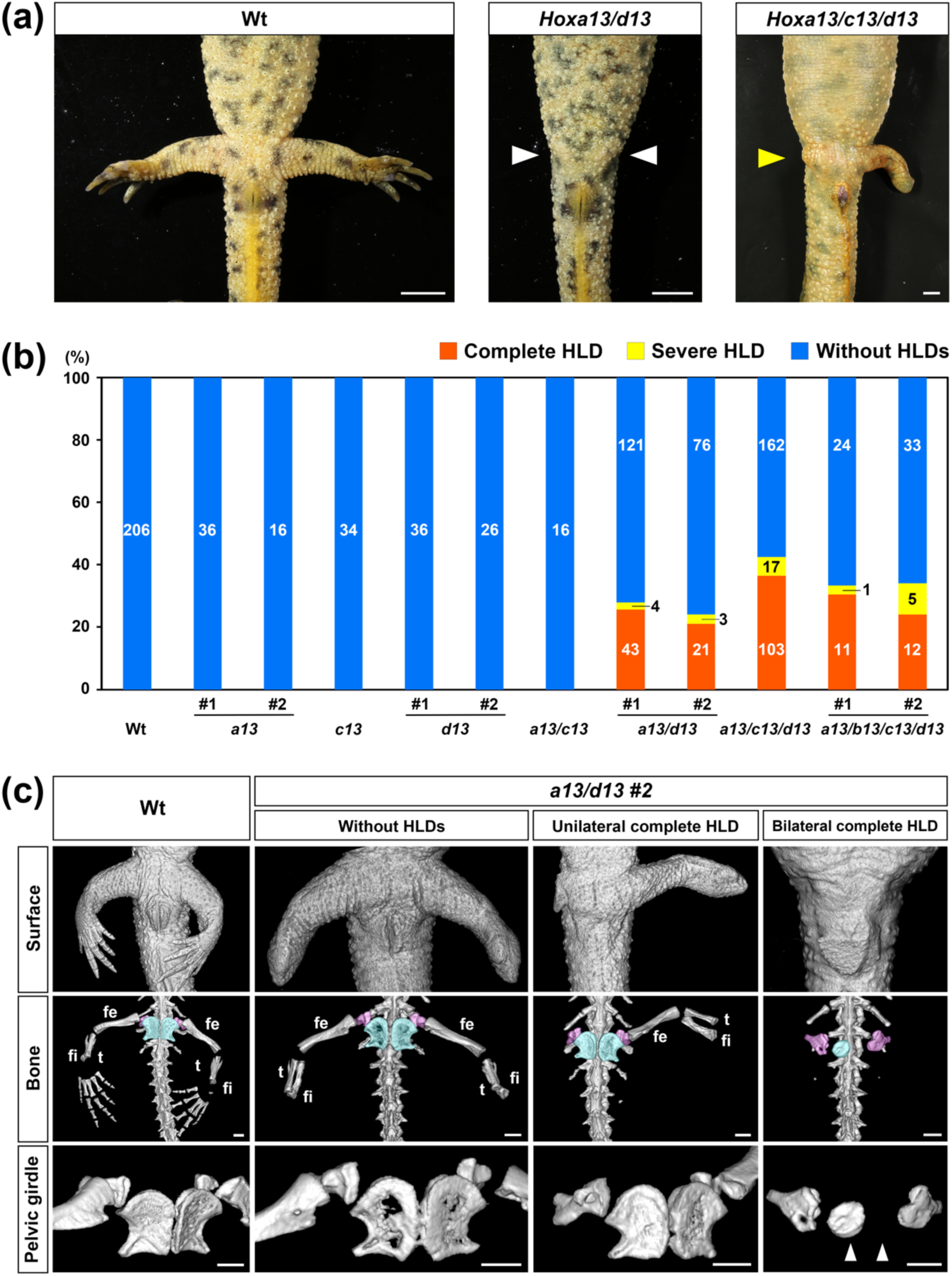
Hindlimb defects in *Hox13* crispants. (a) The left and middle panels are ventral image examples of wild-type newts (Wt) and *Hox a13/d13* crispants with hindlimb defects (HLDs), respectively, at approximately five months post-fertilization (mpf). The right panel is a ventral image example of *Hoxa13/c13/d13* crispant with HLDs at approximately one year post-fertilization. White and yellow arrowheads indicate complete and severe HLD, respectively. Scale bar, 5 mm. (b) Percentages of numbers of the lateral sides with and without HLDs for each genotype. Numbers of sides with each phenotype are shown in the bars. Two crispant groups (#1 and #2) were generated by using different gRNAs (see Table S1). (c) Micro-CT images (ventral) of hindlimbs and pelvic girdles of Wt newts and newts in *Hoxa13/d13* crispant group #2 (*a13/d13 #2*) with complete HLD at around 6 mpf. The middle and lower panels represent whole bones around hindlimbs and pelvic girdles viewed from diagonally right, respectively. Regions shown by light blue and pink colors in the middle panels indicate the ischiopubis and ilium, respectively. White arrowheads in the lower panels indicate pelvic girdle malformation. fe, femur; fi, fibula; t, tibia. Scale bar, 1 mm.

To elucidate how *Hox13* controls whole hindlimb development, we first examined which *Hox13* paralogs were responsible for HLDs by comparing the phenotypes of single, double, triple, and quadruple *Hox13* crispants, which were described in a previous report (Takeuchi et al., 2022). Each *Hox13* paralog gene was targeted by multiple gRNAs (Table S4) (Takeuchi et al., 2022). Two crispant groups (#1 and #2) were analyzed for *Hoxa13*, *Hoxd13*, *Hoxa13/d13*, and *Hoxa13/b13/c13/d13* mutations (Table S4, Figure 1(b)). HLDs were observed in all groups of *Hoxa13/d13* double, *Hoxa13/c13/d13* triple, and *Hoxa13/b13/c13/d13* quadruple crispants (hereafter abbreviated as *a13/d13*, *a13/c13/d13*, and *a13/b13/c13/d13*, respectively), however, not in other crispant groups (Figure 1(b)), indicating that disruption of at least *Hoxa13* and *Hoxd13* was required for the HLDs. As HLDs showing no limbs were observed on either the unilateral or bilateral sides in these crispant groups (Figure 1(a) and 1(c) and Table S5), the percentages of HLDs were calculated using total numbers of lateral sides (Figure 1(b)) or total newts (Table S5) as the baseline. For example, in the case of *a13/d13 #1* crispant group, the percentage per lateral side was 28.0% (47 of 168 sides, Figure 1(b)), and that per total newts was 40.5% (34/ total 84 newts, Table S5). Overall, almost half of *Hox13* crispants showed HLDs (153 newts among 315 *Hox13* crispants, Table S5). The unilateral phenotype appeared more frequently than the bilateral phenotype in these crispants (Table S5). No difference was observed in the frequency of HLDs between the left and right sides in crispants with unilateral HLDs (Table S5).

Next, we analyzed the skeletal morphology of the posterior trunk region in three wild-type newts and 26 *Hox13* crispants (13 newts with HLDs and 13 newts without HLDs), around six months post-fertilization (mpf), using micro-computed X-ray tomography (micro-CT). In total, the 16 lateral sides with HLDs in 13 *Hox13* crispants included 15 sides with complete HLD and one side with severe HLD. All sides with complete HLD had no hindlimb bones (for example, see Figure 1(c)). However, a side with severe HLD had rudimentary femurs (Figure S2(a)). Almost all lateral sides with HLDs in these crispants also showed various pelvic girdle malformations, such as hypoplasia (for example, arrowheads in Figure 1(c); for other malformations, see Figure S2(b)). The frequency of pelvic girdle malformations on the sides with the HLDs (13/16 sides) was significantly higher than that on the sides without HLDs (2/26 sides; P<0.001, Fisher’s exact test). Thus, these results suggest that *Hox13* is involved in the development of whole hindlimbs and pelvic girdles.

### 3.2 Mutation in *Hox13* genes resulted in hindlimb defects

Although genes can be disrupted with extremely high mutation rates by CRISPR/Cas9 in *P. waltl*, these crispants exhibit genetic mosaicism (Suzuki et al., 2018; Takeuchi et al., 2022). Phenotypic variation in HLDs in *Hox13* crispant newts (Figure 1(a)-(c)) might have resulted from this mosaicism. To examine the possibility, we generated *Hox13* germline mutants from the *a13/c13/d13* crispants and analyzed their phenotypes. The pedigree of these germline mutants is described in Material and Methods. We obtained 268 offspring from the four pairs (Table 1). In total, seven mutant newts and 10 sides showed HLDs (nine sides, complete HLD; one side, severe HLD). Detailed genotypes of the mutants with HLDs and their parents and sides with HLDs are shown in Table S2. HLDs appeared in newts with only three genotypes (*a13^-/-^c13^+/-^d13^-/-^*, *a13^-/-^c13^-/-^d13^-/-^*, and *a13^-/-^c13^-/-^d13^+/-^*) (Table 1), suggesting that the HLDs did not result from off-target effects. Similar to crispants, HLDs were observed in a part of the newts with mutations in these three genes (Table 1). In addition, defects were found on either the unilateral or bilateral sides (Figure 2, upper panels and Table S2), showing that phenotypic variation did not result from genetic mosaicism, and occurred in germline mutants with the same genotypes and also on the lateral sides with identical genetic backgrounds. We could not obtain *a13^-/-^c13^+/+^d13^-/-^* newts with HLDs, probably because of the small number of offspring with this genotype (three newts, Table 1).

**FIGURE 2.**
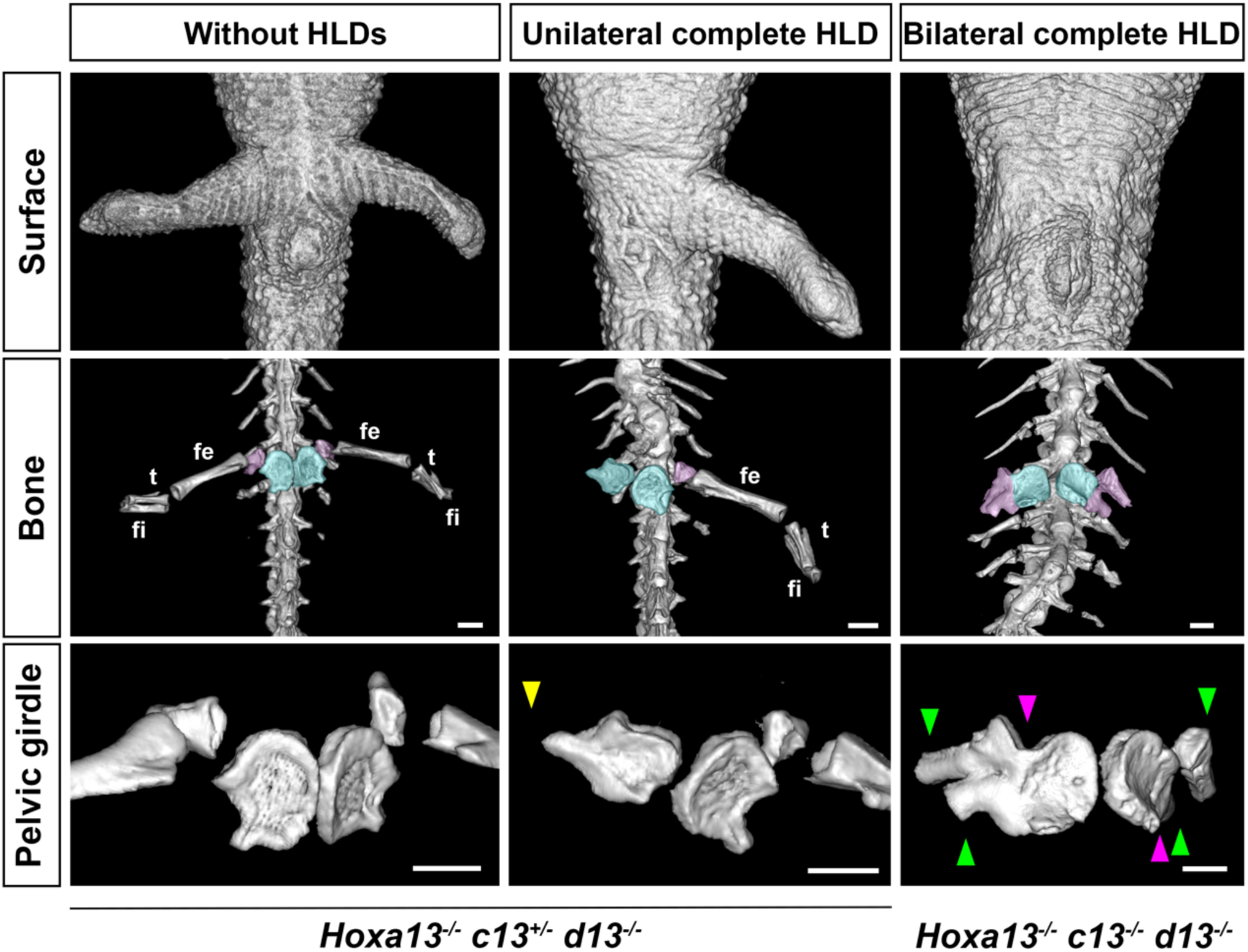
Hindlimb defects in *Hox13* germline mutants. Micro-CT ventral images of hindlimbs and pelvic girdles of *Hox13* germline mutants at approximately 6 mpf. Genotypes are shown at the bottom of the figure. The middle and lower panels represent the entire bone around the hindlimbs and pelvic girdles. The regions shown by colors and abbreviations are the same as those in Figure 1. The yellow, green, and pink arrowheads indicate hypoplasia in the ilium, bifurcation in the ilium, and fusion of the ischiopubis and ilium, respectively. Scale bar, 1 mm.

**Table 1.**
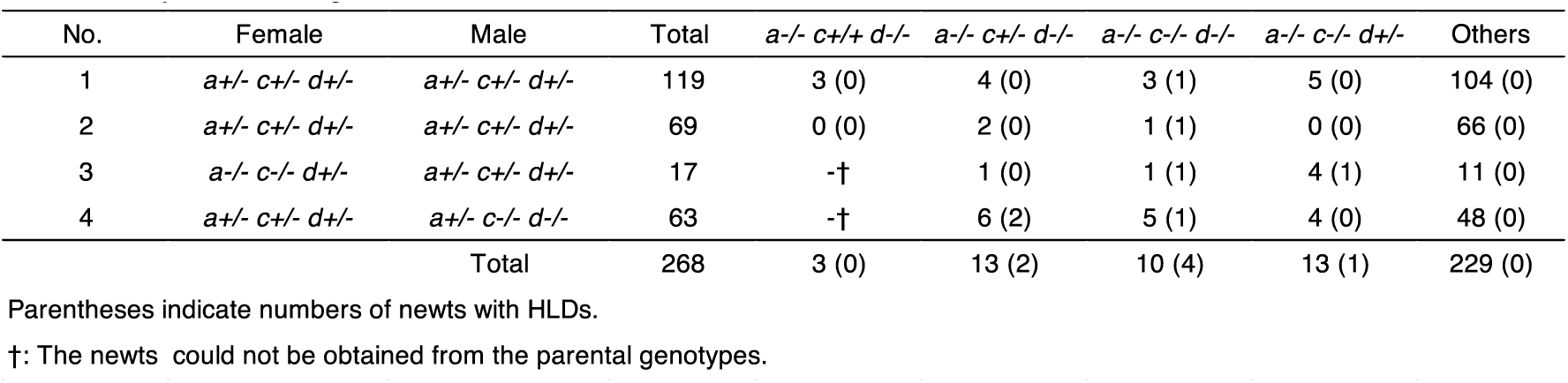
Analysis of *Hox13* germline mutants.

We also analyzed the skeletal morphology of the six lateral sides with HLDs in the *Hox13* germline mutants using micro-CT. Similar to the crispant phenotypes, five sides with complete HLD had no hindlimb bones (for example, Figure 2, middle and bottom panels), whereas one side with severe HLD had only rudimentary femurs (for example, see Figure S2(c)). Five sides with HLDs also showed malformations of the pelvic girdles on the affected side (for example, Figure 2, arrowheads).

Given that *Hox13* crispants and germline mutants showed almost identical phenotypes in HLDs, we used crispants to investigate the functions of *Hox13* genes in the following experiments. Collectively, these results showed that mutations in *Hox13* led to HLDs and that even the identical genotypes and genetic backgrounds caused phenotypic variation.

### 3.3 Hindlimb buds do not undergo outgrowth in *Hox13* crispants

We observed the hindlimb developmental process of *a13/c13/d13* crispants to determine when and how HLDs occurred. We found that the *a13/c13/d13* crispants, which eventually exhibited complete HLD, initially formed limb buds; however, these limb buds did not undergo outgrowth (Figure 3, orange arrowheads). This result was confirmed on all lateral sides with HLDs as far as was examined (35 sides of *a13/c13/d13* crispants and 18 sides of *a13/ d13* crispants). This suggests that *Hox13* is involved in the outgrowth of hindlimb buds in newts. Furthermore, we found that around 20 days post-fertilization (dpf), the morphology and size of the limb buds on one side, which subsequently showed HLDs, were smaller in size with a rounded shape, whereas hindlimb buds on the other side, which did not subsequently show HLDs, were larger with a shape extending in the posterior direction (Figure 3). These differences allowed us to predict HLDs. These results showed that the HLDs of *Hox13* mutant newts resulted from the outgrowth failure of hindlimb buds.

**FIGURE 3.**
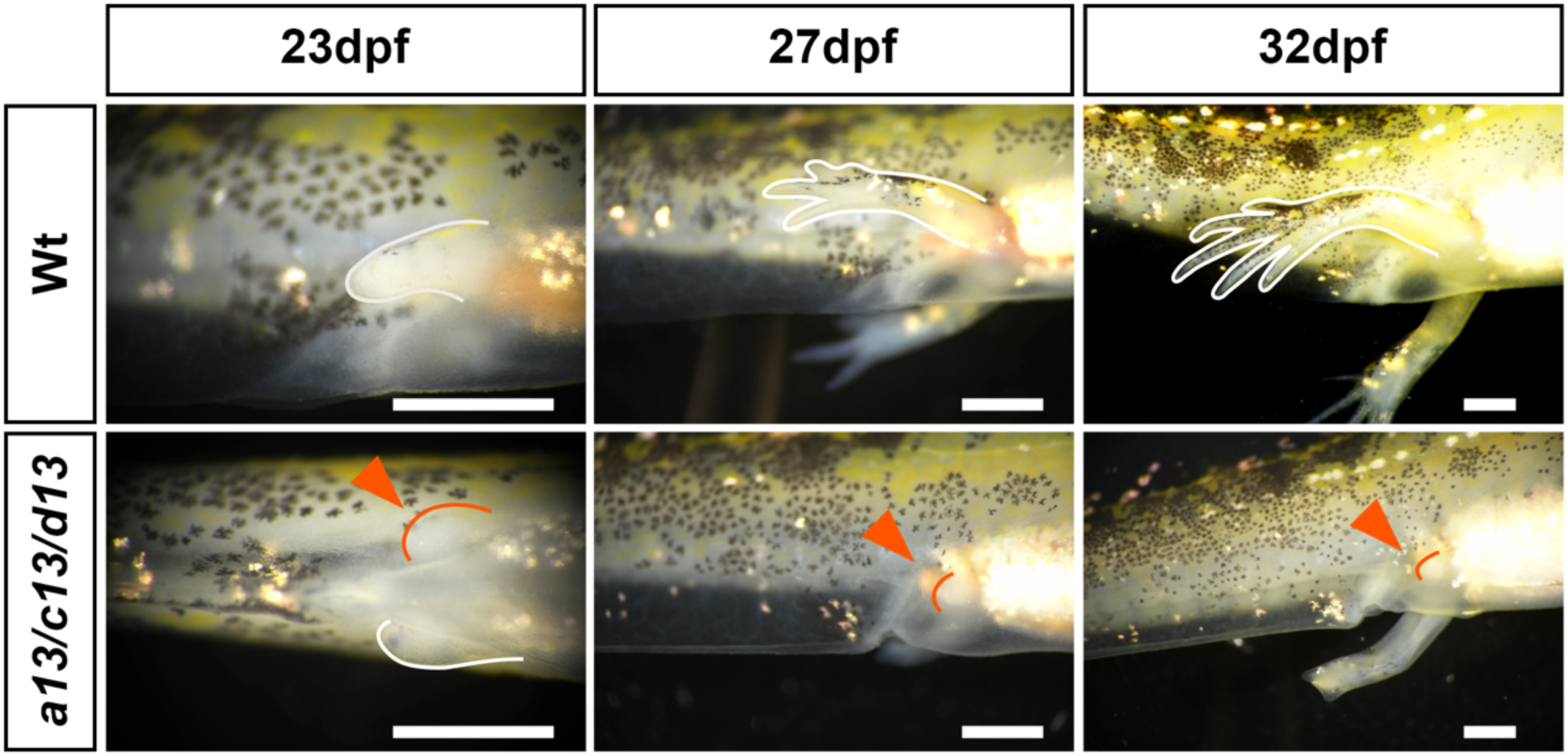
Hindlimb buds did not elongate in *Hox13* crispants. Hindlimb development in a wild-type newt (Wt) and *a Hoxa13/c13/d13* crispant newt (*a13/c13/d13*) is shown. Orange arrowheads show the hindlimb with complete hindlimb defect. Scale bar, 1 mm.

### 3.4 Newt *Hox13* regulates the expression of genes controlling hindlimb development

We examined the expression patterns of *Hoxa13*, *Hoxd13, Fgf10, Tbx4* and *Pitx1* during hindlimb development using WISH. For all genes, antisense probe-specific signals were detected in the hindlimb buds of wild-type newts (Figure S3). The expression of *Hoxa13* was detected in the distal regions of wild-type newts at the middle stage (around 20dpf) but not in the early stage (around 18 dpf), although the signal was very weak (Figure 4(a), dotted lines). The expression signal became stronger at the late stages (around 22 dpf) (Figure 4(a)). In contrast, *Hoxd13* expression was detected only in the posterior-distal region at the late stage (Figure 4(a)). These results showed that the expression patterns of *Hoxa13* and *Hoxd13* are similar in newt forelimbs and hindlimbs (Takeuchi et al., 2022).

**FIGURE 4.**
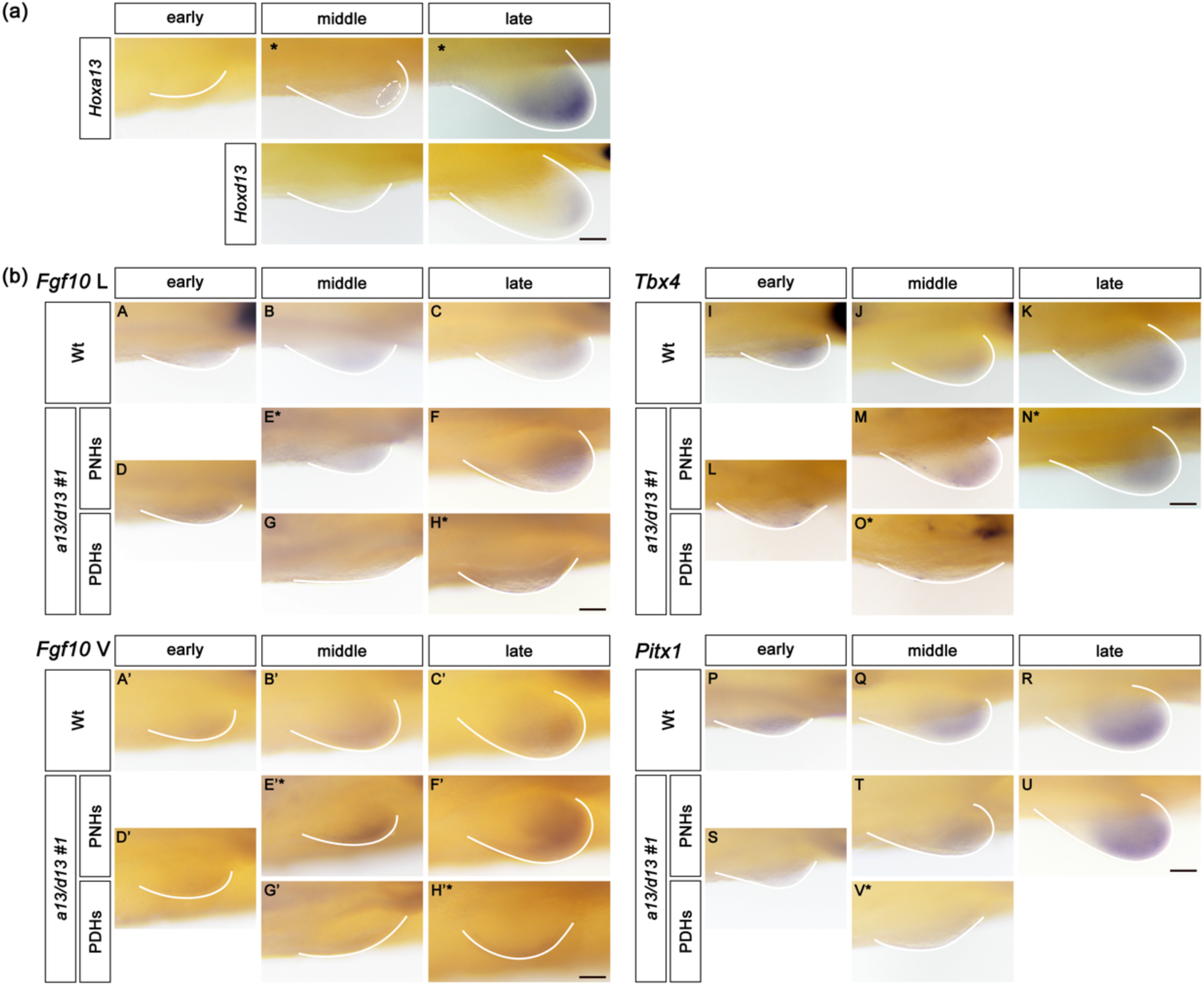
Newt *Hox13* regulates the expression of genes controlling hindlimb development. Gene expression patterns were analyzed using whole-mount in situ hybridization. Asterisks indicate images of the left hindlimbs reversed to align with the orientation of the right hindlimbs. (a) *Hoxa13* and *Hoxd13* expression pattern in the hindlimb buds of wild-type newts. The area surrounded by the dotted line indicates a weak expression region of *Hoxa13.* Scale bar, 0.1 mm. (b) Expression patterns of *Fgf10, Tbx4*, and *Pitx1*, in hindlimbs of wild-type and *Hox13* crispant newts. Photographs other than A’-H’ represent lateral views. The same hindlimbs were photographed from the lateral (*Fgf10* L: A-H) and ventral sides (*Fgf10* V: A’-H’). For example, A and A’ are the same hindlimb. Wt: wild-type newts, PNHs: predicted normal hindlimbs regarding the outgrowth, PDHs: the predicted defective hindlimbs. “early,” “middle,” and “late” indicate limb developmental stages (around 18 dpf, 20 dpf, and 22 dpf, respectively). E and G, F and H, M and O, T and V are the hindlimbs of the same newts predicted to have unilateral defects. Scale bar, 0.1 mm.

Next, we compared the expression patterns of *Fgf10*, which is involved in limb field specification, limb bud formation, and outgrowth (Min et al., 1998; Ohuchi et al., 1997; Sekine et al., 1999), between wild-type and *Hox13* crispant newts. We predicted hindlimb buds with defective and normal outgrowth (hereafter, PDHs and PNHs, respectively) based on their morphology from the middle to late stages (Figure 3). *Fgf10* expression was broadly detected in the hindlimbs of wild-type newts, even at an early stage (Figure 4(b)A), suggesting that *Fgf10* was expressed before the start of *Hox13* expression. No apparent differences were observed in *Fgf10* expression at the early stage between the wild-type and *Hox13* crispant newts (Figure 4(b), A and D, A’ and D’), although PDHs and PNHs could not be distinguished. However, *Fgf10* expression was markedly reduced in PDHs at the middle and late stages compared to that in PNHs and wild-type hindlimb buds (numbers of sides analyzed: PDHs, three sides; PNHs, 17 sides; wild-type, 12 sides) (Figure 4(b), E* and G, F and H*). These results suggested that *Fgf10* was downregulated after the middle stage in the limb buds on the HLD side of *Hox13* crispant newts.

Additionally, we analyzed the expression of *Tbx4* and *Pitx1* genes. *Tbx4* regulates hindlimb development as a gene upstream of *Fgf10* (Naiche & Papaioannou, 2003). *Pitx1* determines hindlimb identity and regulates *Tbx4* expression (Logan & Tabin, 1999). The spatiotemporal expression patterns of these two genes were similar to those of *Fgf10* (Figure 4(b), A-C, I-K and P-R). Their expression seemed to be slightly reduced in *Hox13* crispants compared to wild-type newts at the early stages (Figure 4(b), I and L, P and S), although PDHs and PNHs could not be distinguished. At the middle stages, their expression was strongly reduced in PDHs compared with PNHs (Figure 4(b), M and O*, T and V*) (numbers of sides analyzed: for *Tbx4*, PDHs, five sides, PNHs, seven sides, wild-type, eight sides; and for *Pitx1,* PDHs, three sides, PNHs, 15 sides, wild-type, 24 sides). These results suggested that *Tbx4* and *Pitx1* were also downregulated after the middle stage in limb buds on the HLDs side of *Hox13* crispants. The stages at which *Fgf10, Tbx4,* and *Pitx1* expression was reduced are consistent with the stages at which *Hox13* expression started. These results suggested that *Hox13* regulated the expression of these three genes during hindlimb development.

### 3.5 Mutations in *Fgf10* and *Tbx4*, however, not in *Pitx1* cause complete hindlimb defect

As *Fgf10, Tbx4,* and *Pitx1* expression decreased in *Hox13* crispants (Figure 4), *Hox13* might have regulated hindlimb development by interacting with these two genes. Therefore, we mutated these genes using the CRISPR-Cas9 system and compared the resulting phenotypes with those of *Hox13* crispants. gRNAs with high mutation rates in *P. waltl* (Suzuki et al., 2024) were used for *Fgf10* (Table S6). The target regions of *Tbx4* and *Pitx1* gRNAs were designed with reference to mouse and zebrafish mutations (Table S6) (Don et al., 2016; Lanctôt, Moreau, Chamberland, Tremblay, & Drouin, 1999; Naiche & Papaioannou, 2003).

Amplicon sequencing was performed for genotyping by next-generation sequencing (NGS). The genomic DNAs from two to seven newts in each group were investigated. All target sites for *Fgf10* and *Tbx4* crispants were mutated with high efficiency (86.5– 97.2%) (Table S7). In contrast, the *Pitx1* crispants were divided into two groups. One group (four newts) showed high mutation efficiency (mean, 90.6%) and abnormal phenotypes, as described below, whereas the other group (three newts) showed low mutation efficiency (mean, 40.7%) and no abnormal phenotypes (Table S7). Therefore, only crispants showing abnormal phenotypes were examined in subsequent analyses. Mutations in all gRNAs of *Fgf10, Tbx4*, and *Pitx1* yielded deletions or insertions around target regions, primarily causing frameshift mutations (Table S8), suggesting that the functions of the target genes were disrupted.

A part of *Fgf10* mutant newts exhibit fewer digits and the absence or hypoplasia of zeugopods in the hindlimbs (Suzuki et al., 2024). We generated *Fgf10* crispants using the same gRNAs and confirmed almost the same phenotypes (at least, fewer digits) in 89.7% (gRNA #1) and 67.9% (gRNA #2) of *Fgf10* crispants (Table S9; for examples, Figure 5a). In addition, we found complete HLD in a few sides (gRNA #1: 3.8%, n=78 sides; #2: 2.2%, n=134 sides; Table S9) (for examples, Figure 5(a)). We observed the developmental process of the hindlimb in *Fgf10* crispants and found that all lateral sides of the *Fgf10* crispants formed hindlimb buds. This suggests that the defect was related to hindlimb bud outgrowth. Complete HLD was observed on six sides (one bilateral and four unilateral). In addition, two sides of one crispant showed severe HLD (Figure S4(a)). Both complete and severe HLD were similar to those observed in *Hox13* crispants (Figures 1(c) and S2(a)). Almost all *Fgf10* crispants died around the metamorphosis stages for unknown reasons. We also examined *Fgf10* germline mutant (*Fgf10*^-/-^) newts (Suzuki et al., 2024). F0 animals were generated using the same gRNA as gRNA #2 (Tables S1 and S6, G78). Complete and severe HLD were observed in seven sides (20.6%) and five sides (14.7%) of the 34 sides of 17 *Fgf10*-/- newts, respectively (for examples, Figure S4(b)).

**FIGURE 5.**
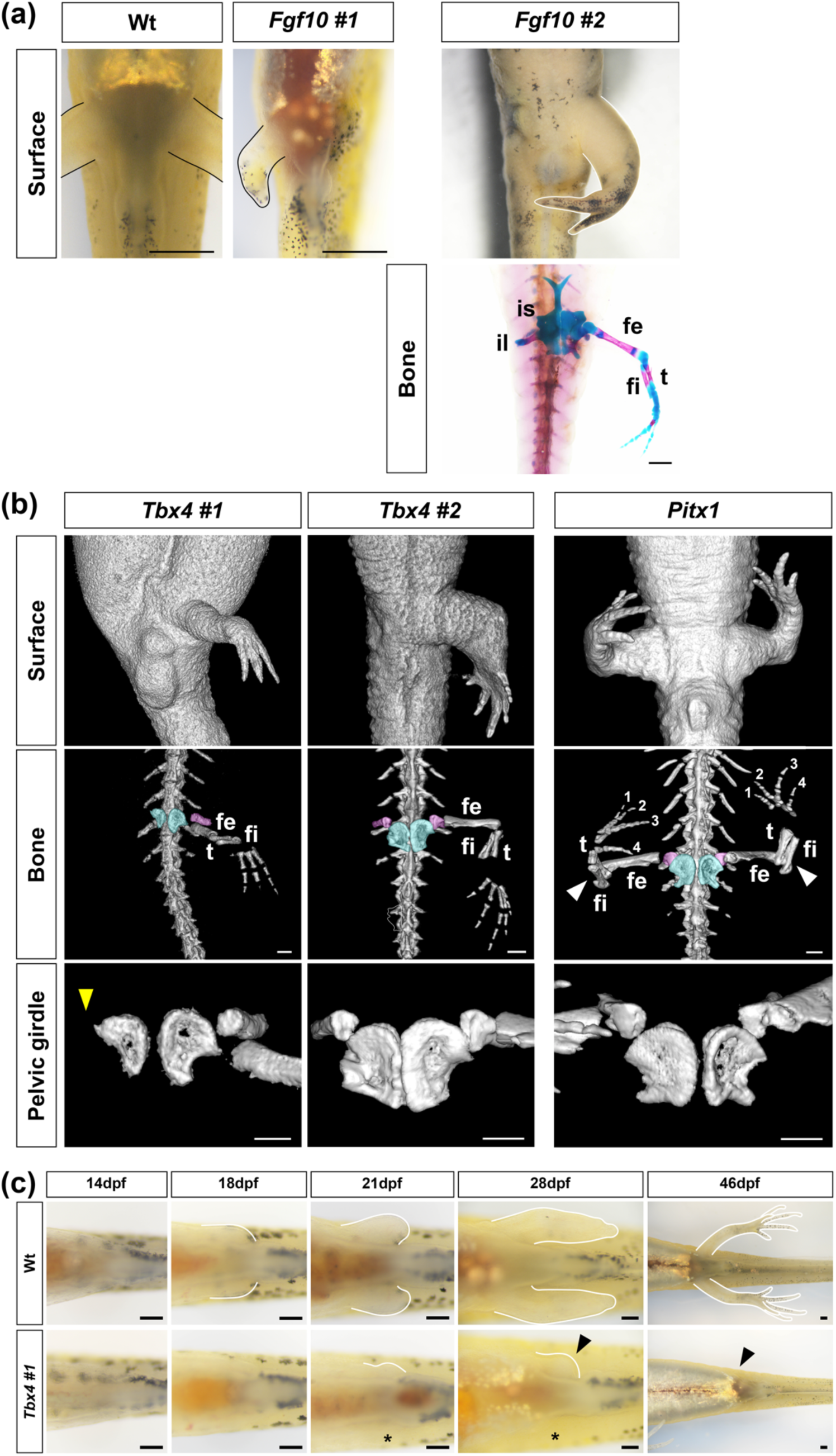
Hindlimbs and pelvic girdles in *Pitx1, Tbx4,* and *Fgf10* crispants. (a) Ventral views of the hindlimbs with wild-type (2mpf), *Fgf10* guide RNA #1 (2mpf), and *Fgf10* guide RNA #2 (approximately 4 mpf) crispant newts. The bone and cartilage staining image is also shown for a *Fgf10* guide RNA #2 crispant. Abbreviations are the same as those in Figure 1. Scale bar, 1 mm. (b) Micro-CT images (ventral) with *Pitx1*, *Tbx4* guide RNA *#1*, and *Tbx4* guide RNA *#2* crispant newts at 5–6 mpf. The regions shown by colors and abbreviations are the same as those in Figure 1. The yellow and white arrowheads indicate hypoplasia in the ilium and anterior orientation of the stylopods and zeugopods. Scale bar, 1 mm. (c) Hindlimb development of wild-type and *Tbx4* guide RNA #1 crispant newts. Ventral views. Arrowheads: hindlimb buds at a LB + side, in which a hindlimb bud was formed, Asterisks: a LB – side, in which a hindlimb bud was not formed. Both sides eventually showed complete hindlimb defects. Scale bar, 0.2 mm.

Next, we examined the hindlimbs of the *Tbx4* crispants. Many *Tbx4* crispants died after metamorphosis for unknown reasons in both crispant groups (#1 and #2). Most *Tbx4* crispants exhibited complete HLD (gRNA #1: 54.5%, n=112 sides; #2: 78.3%, n=230 sides) (for examples, Figure 5(b)) (Table 2). No case of severe HLD was observed. Complete HLD was observed on unilateral and bilateral sides (19.5% and 80.5%, respectively, in 133 crispants with complete HLD). As similar phenotypes were observed in crispants with both gRNAs, subsequent analyses were performed using gRNA #1. We examined hindlimb bud formation in *Tbx4* crispants around 24 dpf, and found that 70 out of 112 sides (62.5%) formed limb buds (hereafter referred to as LB+ sides) (Table 2, for example, Figure 5(c), arrowheads), whereas the remaining 42 sides (37.5%) did not form the limb buds (hereafter referred to as LB- sides) (Table 2, for example, Figure 5(c), asterisks). As all *Hox13* crispants formed limb buds as far as we examined, we first focused on LB+ sides of *Tbx4* crispant newts. Complete HLD was observed in 23 of the 70 LB+ sides (32.9%) (Table 2, for example, see Figure 5(c), arrowheads), suggesting a failure in hindlimb bud outgrowth. The morphology of these limb buds was similar to that observed in *Hox13* crispants (arrowheads in Figures 3 and 5(c)). These results suggested that *Tbx4* was involved in the outgrowth of hindlimb buds.

**Table 2.** Hindlimb phenotypes of *Tbx4* gRNA *#1* crispant newts.

|  | Without HLDs† | Short | Complete HLD† | NE‡ | Total |
| --- | --- | --- | --- | --- | --- |
| LB+ sides§ | 39 (55.7) | 6 (8.6) | 23 (32.9) | 2 (2.9) | 70 (100) |
| LB- sides¶ | 2 (4.8) | 0 (0) | 38 (90.5) | 2 (4.8) | 42 (100) |

Parentheses indicate the percentages that amount of the LB+ sides§ or LB- sides¶.
† HLD(s): hindlimb defect(s)
‡ NE: not examined
§LB+ sides: Sides in which hindlimb buds were formed.
¶LB- sides: Sides in which hindlimb buds were not formed.

Short hindlimbs were also found on the LB+ sides (8.6%, Table 2; hereafter referred to as short HLD). As these limbs mainly lacked upper legs (Figure S5, short HLD), the phenotype differed from that of severe HLD in the *Hox13* crispants, which lacked entire lower legs and feet and had only rudimentary femurs (for example, Figure S2(a)). Almost all LB- sides (90.5%) showed complete HLD (Table 2), suggesting that newt *Tbx4* also specified the limb field or limb bud formation within the field. This is a novel result and is different from the *Tbx4* mutant zebrafish and mice, which form fin buds and limb buds (Don, Hall, Currie, & Cole, 2011; Naiche & Papaioannou, 2007). Pelvic abnormalities were observed in *Tbx4* crispants (Figure 5(b), yellow arrowhead). Some crispants were missing all the pelvic girdles (Figure S5). Whether these abnormalities were present on the LB+ or LB- side was not examined in all cases.

Almost half of the *Pitx1* crispants showed abnormal phenotypes (31 out of 64 newts, 48.4%). As described above, crispants without abnormal phenotypes showed low mutation efficiencies (Table S7); therefore, only crispants showing abnormal phenotypes were subsequently examined. Unlike *Hox13, Fgf10*, and *Tbx4* crispants, no *Pitx1* crispants had complete HLD. Instead, these crispants exhibited both of the following two phenotypes in the hindlimbs. Digit numbers were reduced from five to four, and the stylopod and zeugopod oriented anteriorly similar to forelimbs (white arrowheads in Figure 5(b)). These abnormalities were observed bilaterally in almost all newts with abnormal phenotypes (30 out of 31 newts). No apparent pelvic girdle malformations were observed. Thus, *Pitx1* crispants did not show the same phenotypes as *Hox13* crispants.

These results show that *Fgf10* and *Tbx4* regulated the outgrowth of hindlimb buds and support the possibility that *Hox13* regulates outgrowth by interacting with *Fgf10* and *Tbx4*.

### 3.6 *Hox13* has genetic interactions with *Fgf10* and *Tbx4*

*Fgf10* and *Tbx4* crispants exhibited phenotypes similar to those of *Hox13* crispants, such as complete HLD (Figure 5). To examine the genetic interaction between *Hox13* and *Fgf10* or *Hox13* and *Tbx4*, we generated compound crispants (*Hoxa13/d13*/*Fgf10* or *Hoxa13/d13*/*Tbx4*) and compared their phenotypes with those of the original crispants (*Hoxa13/d13, Tbx4* or *Fgf10* crispants). gRNAs for *Hoxa13/d13 #1, Fgf10 #1* and *Tbx4 #1* were used (Tables S1 and S5).

All lateral sides of *Hoxa13/d13/Fgf10* crispants formed hindlimb buds. The percentage of complete HLD in *Hoxa13/d13*/*Fgf10* crispant sides (70.8%, n=48) significantly increased compared to that in the original crispant sides (Figure 6(a)) (*Hoxa13/d13*: 25.6%, n=168; *Fgf10*: 3.8%, n=78). Complete HLD of *Hoxa13/d13*/*Tbx4*, *Hoxa13/d13* and *Tbx4* crispants were compared. The percentage of complete HLD in *Hoxa13/d13*/*Tbx4* LB+ sides (54.9%, n=51) was significantly higher than that in *Hoxa13/d13* (25.6%, n=168) or *Tbx4* LB+ sides (32.9%, n=70; same data in Table. 2) (Figure 6(b)). These results indicate that *Fgf10* and *Tbx4* genetically interacted with *Hox13*. Collectively, our results show that *Hox13* regulated the outgrowth of hindlimb buds by interacting with *Fgf10* and *Tbx4*.

**FIGURE 6.**
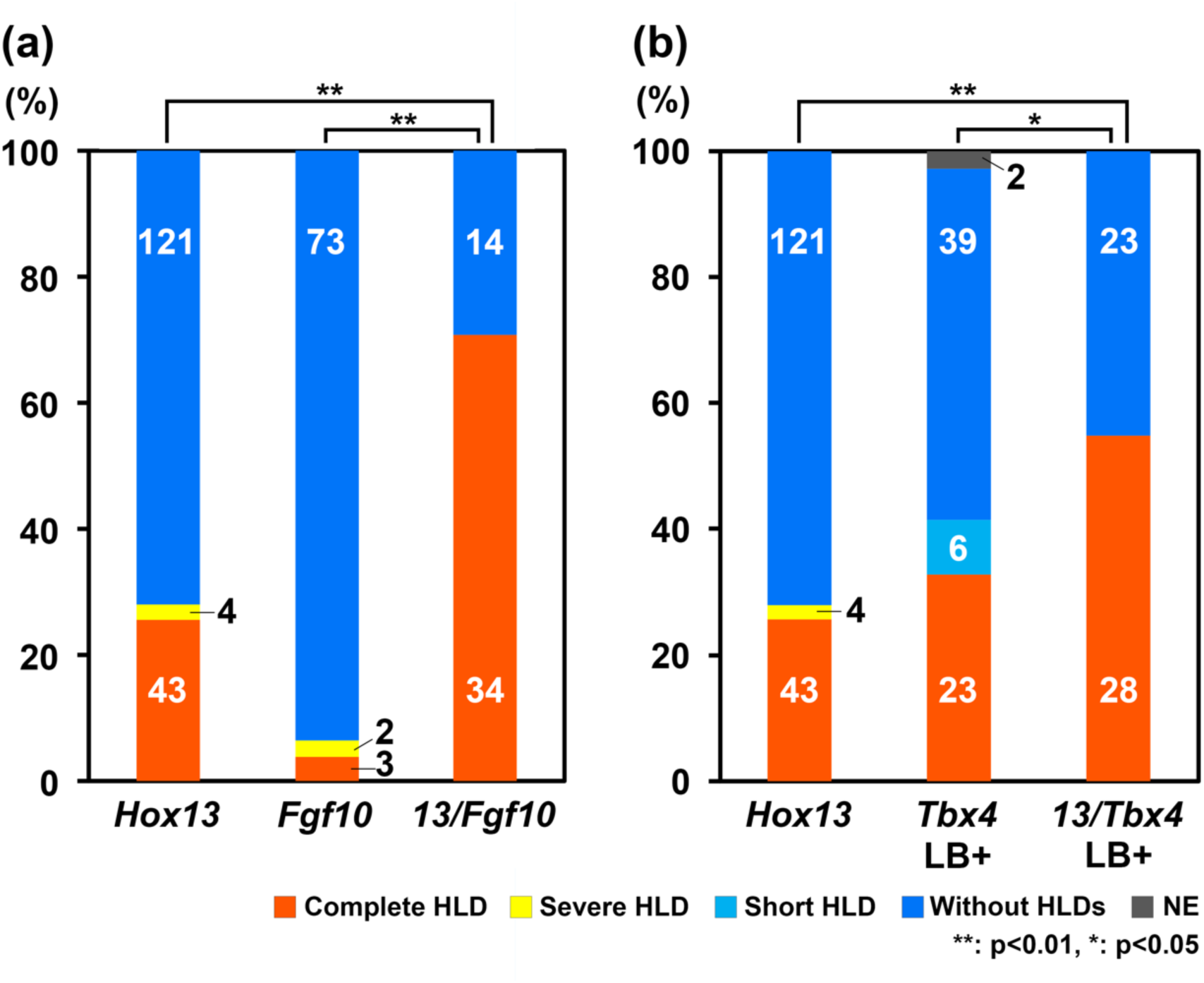
*Hox13* had genetic interactions with *Fgf10* and *Tbx4*. The percentages and numbers of lateral sides for each phenotype are shown. (a) Data for *Hox13* (*Hoxa13/d13 #1*), *Fgf10* (*Fgf10 #1*), and the compound crispant *13/Fgf10* (*Hoxa13/d13 #1/Fgf10 #1*) sides. (b) Data for *Hox13* (*Hoxa13/d13 #1*), *Tbx4* LB+ (*Tbx4 #1*; same data in Table 2), and the compound crispant *13/Tbx4* (*Hoxa13/d13 #1/Tbx4 #1*) LB+ sides, in which hindlimb buds were formed. HLDs: hindlimb defects, NE: not examined. *: p<0.05, **: P<0.001, the significance was tested for differences in the number of sides with or without complete HLD (Fisher’s exact tests).

## 4 DISCUSSION

### 4.1 *Hox13* regulates outgrowth of hindlimb buds

*Hox13* genes regulate the formation of digits in mice (Fromental-Ramain, Warot, Messadecq, et al., 1996) and newts (Takeuchi et al., 2022) and fin rays in zebrafish (Nakamura, Gehrke, Lemberg, Szymaszek, & Shubin, 2016). However, the present study is the first to report that the newt *Hox13* gene regulated whole hindlimb (posterior appendages) development. All sides with HLDs in *Hox13* crispant newts had hindlimb buds; however, the buds did not show outgrowth (Figure 3). *Hox13* expression was detected first at the middle stage in limb buds, though not at the early stages (Figure 4(a)). These results suggested that *Hox13* regulated the outgrowth of newt hindlimb buds.

### 4.2 Putative *Hox13*-mediated regulation of hindlimb bud outgrowth

We showed that the expression of *Fgf10* and *Tbx4*, which are involved in limb development, was markedly reduced in the PDHs of *Hox13* crispants (Figure 4(b)). In addition, the hindlimb buds did not show outgrowth in a part of either *Fgf10* or *Tbx4* crispants. *Hox13* promoted the outgrowth of hindlimb buds by interacting with *Fgf10* and *Tbx4* (Figure 6). These results suggest that *Hox13* regulated the outgrowth of hindlimb buds by controlling *Fgf10* and *Tbx4* expression*. Fgf10* expression decreased in the forelimbs of *Hox13* knock-out mice, and Hox13 proteins bound to the franking regions of *Fgf10* gene (Sheth et al., 2016), suggesting that Hox13 directly controls *Fgf10* expression. Although, whether Hox13 directly regulates *Tbx4* expression remains unknown, *Tbx4* regulates hindlimb outgrowth through the activation of *Fgf10* in mice (Naiche & Papaioannou, 2003, 2007). Based on these reports and the present data, we hypothesized a novel *Hox13* cascade. In this *Hox13/Tbx4*/*Fgf10* cascade, *Hox13* activates both *Tbx4* and *Fgf10,* whereas *Tbx4* activates *Fgf10.* Ultimately, both *Tbx4* and *Fgf10* promote the outgrowth of hindlimb buds (Figure 7(a), *Hox13-*dependent cascade). We believe that the *Hox13/Tbx4*/*Fgf10* cascade works in the distal region of the hindlimb buds from the middle stage based on the expression patterns of *Hox13*, *Tbx4,* and *Fgf10* (Figure 4). This cascade may work before the formation of the hindlimb proximo-distal pattern, which may explain why HLDs can occur in newts.

**FIGURE 7.**
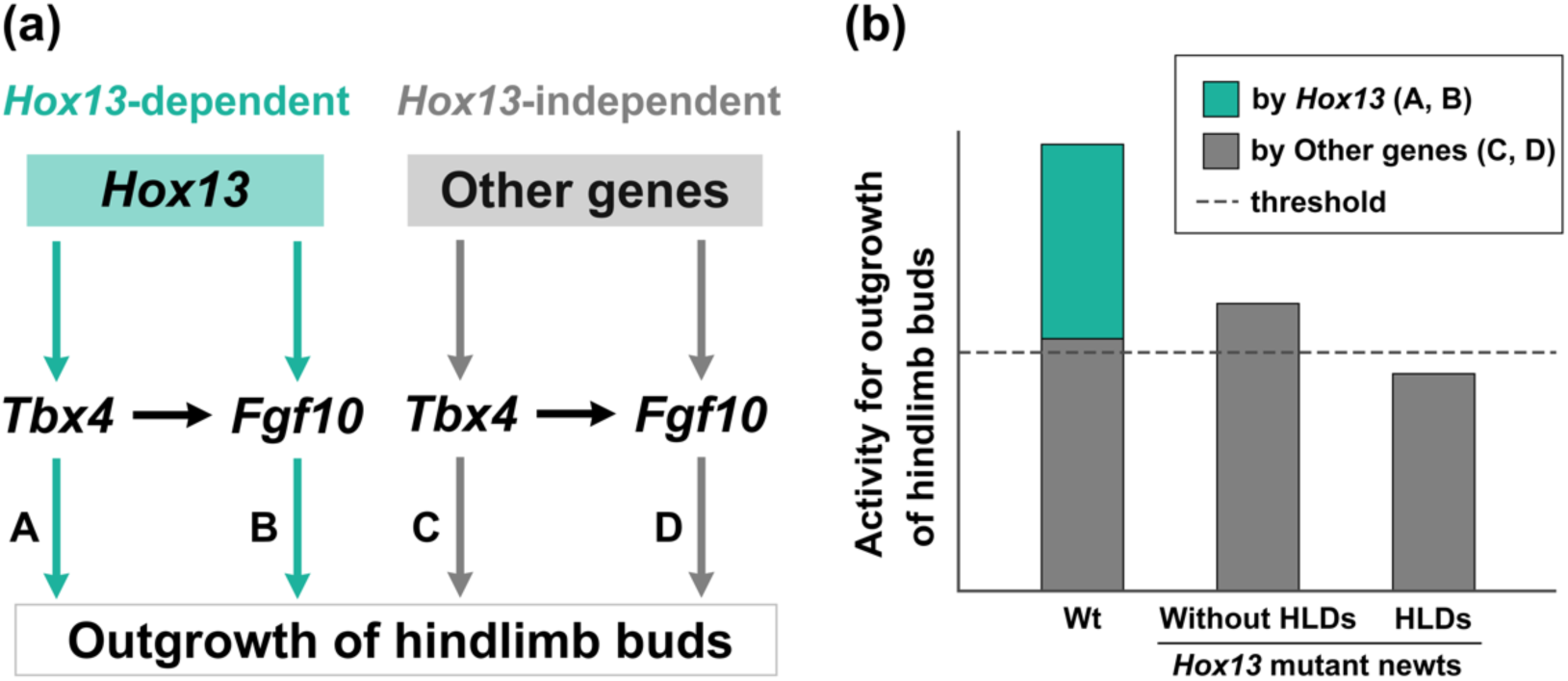
Proposed cascade and model. (a) Proposed cascade. In the *Hox13* dependent cascade, *Hox13* activates both *Tbx4* and *Fgf10,* and *Tbx4* also activates *Fgf10.* Finally, both *Tbx4* and *Fgf10* promote the outgrowth of hindlimb buds. In the *Hox13* independent cascade, genes other than *Hox13* activate both *Tbx4* and *Fgf10. Hox13*-dependent *Tbx4* and *Fgf10* activities on outgrowth of hindlimb buds are shown as A and B and *Hox13*-independent *Tbx4* and *Fgf10* activities as C and D, respectively. (b) Proposed model explaining the difference between the lateral sides of *Hox13* mutants with and without hindlimb defects (HLDs). Achieving a certain threshold level of the combined activities of *Hox13* dependent and independent cascades (wild-type newts, Wt) or only of *Hox13* independent cascade (*Hox13* mutant newts) is crucial for successful outgrowth. The activity of the *Hox13-*independent cascade probably differs between individuals or the lateral sides in *Hox13* mutant newts. The sides with activity higher than the threshold would form the hindlimbs (without HLDs), whereas those with lower activity would not, in the *Hox13* mutant newts (HLDs).

### 4.3 Difference between the lateral sides of *Hox13* mutants with and without HLDs

The factor determining the difference between the lateral sides of *Hox13* mutants with and without HLDs remains unknown. We propose a model based on our results to explain this difference (Figure 7 (b)). We assumed two key points for the model: (1) there is also a *Hox13* independent cascade, in which genes other than *Hox13* promote outgrowth of hindlimb buds through activation of *Tbx4* and/or *Fgf10* (Figure 7(a), *Hox13* independent cascade); and (2) achieving a certain threshold level for the combined activities for outgrowth of hindlimb buds of these two cascades or only a *Hox13-* independent cascade is crucial for successful hindlimb bud outgrowth (Figure 7(b)). The activity of the *Hox13-*independent cascade may differ between individuals or the lateral sides in *Hox13* mutant newts. The sides with higher activity in the *Hox13* mutant newts than the threshold would form the hindlimbs (Figure 7(b), Without HLDs), whereas those with lower activity would not form (Figure 7(b), HLDs).

### 4.4 Difference between newts and mice

Although we observed HLDs in the hindlimbs of *Hox13* mutant newts, HLDs have not been reported in the hindlimbs of mouse *Hox13* mutants (Fromental-Ramain, Warot, Messadecq, et al., 1996). Other redundant genes may compensate for the *Hox13* function that regulates the outgrowth of limb buds in *Hox13* mutant mice. Genes, such as *Shh, Pitx1/2*, and *Islet1*might be involved in the outgrowth. Mice with loss of the limb-specific enhancer (ZRS) of *Shh* had limb buds, although the limbs were markedly reduced (Kvon et al., 2016), suggesting failed outgrowth of limb buds. *Pitx1/2* compound mutant mice showed a reduced hindlimb bud size and the digit 1, tibia, and femur were affected in hindlimbs (Marcil, Dumontier, Chamberland, Camper, & Drouin, 2003). The most severe phenotype (loss of whole hindlimbs) of conditional knockout mice, in which *Islet1* was inactivated in the hindlimb-forming region, suggests that *Islet1* is involved in the initiation of outgrowth of hindlimb buds (Itou et al., 2012). To examine the possibility that other redundant genes compensate for the *Hox13* function in mouse limbs, it would be important to compare the expression of these candidate genes between *Hox13* mutant mice and newts, and analyze compound mouse mutants of *Hox13* and single or multiple of these genes.

### 4.5 Difference between newt forelimbs and hindlimbs

In addition, the loss of whole limbs was not observed in the forelimbs of newt *Hox13* mutants (Takeuchi et al., 2022). Similar to the difference between newts and mice described above, redundant genes may compensate for the *Hox13* function in forelimb buds of *Hox13* mutant newt. *Tbx5* is expressed in the forelimb regions whereas *Tbx4* in the hindlimb regions (for reviews, see Sheeba and Logan, 2017). *Tbx5* expression in the presumptive forelimb region is regulated by *Hox4* and *Hox5* (Minguillon et al., 2012). *Tbx5* plays an essential role in the initiation of limb outgrowth in mice (Agarwal et al., 2003; Rallis et al., 2003) and in newts (Suzuki et al., 2018). These reports suggest that the developmental mechanisms regulated by *Hox4*/*5* and *Tbx5* compensate for *Hox13* function in the forelimb buds of *Hox13* mutant newt.

### 4.6 Insight into hindlimb loss during vertebrate evolution based on the present study

During evolution, various vertebrates have lost their hindlimbs (posterior appendages), probably because of mutations in a variety of genes (for reviews, see Don, Currie, & Cole, 2013; Swank, Sanger, & Stuart, 2021). Such animals might gradually reduce their hindlimb structure and finally lose their entire hindlimbs. Although no known cases of hindlimb loss owing to mutations in *Hox13* have been reported, a PCR survey could not identify the *Hoxd13* gene in Siren, a salamander that completely lost its hindlimbs (Mannaert, Roelants, Bossuyt, & Leyns, 2006). Instead, Siren may have a retro-pseudogene of *Hox13* (Mannaert et al., 2006). However, these require further validation using more detailed analyses, such as whole genome sequencing. *Hoxd9* expression was not detected in pufferfish, which do not have pelvic fins, suggesting the possible role of *Hox* in the loss of the posterior appendages. Notably, the pelvic fin buds were formed, however, did not show the outgrowth (Tanaka et al., 2005), which is similar to the phenotypes observed in *Hox13* mutant newts (Figure 3). Seahorses also lack pelvic fins and the *Tbx4* gene has been deleted from their genome (Lin et al., 2016). In addition, the *Tbx4* mutant zebrafish formed fin buds, although the buds do not show outgrowth and lack pelvic fins (Lin et al., 2016).

Our study suggests that *Hox13* regulates *Tbx4* and *Fgf10* expression. It is possible that the *Hox13/Tbx4/Fgf10* cascade is involved in outgrowth-specific regulation in the posterior appendages and may play a role in the gradual reduction and complete loss of posterior appendages during vertebrate evolution. Further investigation into the role of *Hox13* as a regulator of *Fgf10* and *Tbx4* in hindlimb loss during evolution is required.

Our study revealed that the functions of *Hox13* and the mechanism of hindlimb formation related to *Hox13* are quite different between newts and mice. These results suggest that *Hox* gene functions in vertebrates is more diverse than previously thought. This diversification of *Hox* gene function is expected to correlate with changes in the morphology and function of various tissues across taxa.. Genetic analyses of *Hox* genes in various vertebrates are necessary to test this hypothesis.

## AUTHOR ACONTRIBUTION

ST and TT designed experiments. ST, FM, KS, TH and TT produced the newt crispants and germline mutants. YK, MS and MA performed micro-CT analysis. KS, MM, SS and TH provides the newt genome information. ST and TT performed the other experiments and analyzed the data. HM guided WISH experiments. HM and GA discussed with ST and TT, and contributed to designs and performance of the experiments. ST and TT prepared the manuscript. ST, HM, GA and TT reviewed and edited the manuscript.

## ACKNOWLEDGMENTS

The authors wish to thank Ms. Arisa Anami, Mr. Reo Soeta, Ms. Akiko Adachi and Mr. Hiroshi Onishi (Tottori University, Yonago, Japan) for their technical assistance. This work was supported by JSPS KAKENHI Grant Number JP23KJ1596 to ST. This work was also supported by JSPS KAKENHI JP20K06656 and 23K05783 to T.T. and NIBB Collaborative Research Projects (24NIBB525, 23NIBB201, 23NIBB503, 22NIBB201 and 22NIBB504) to TT.

---

**Table S1.**
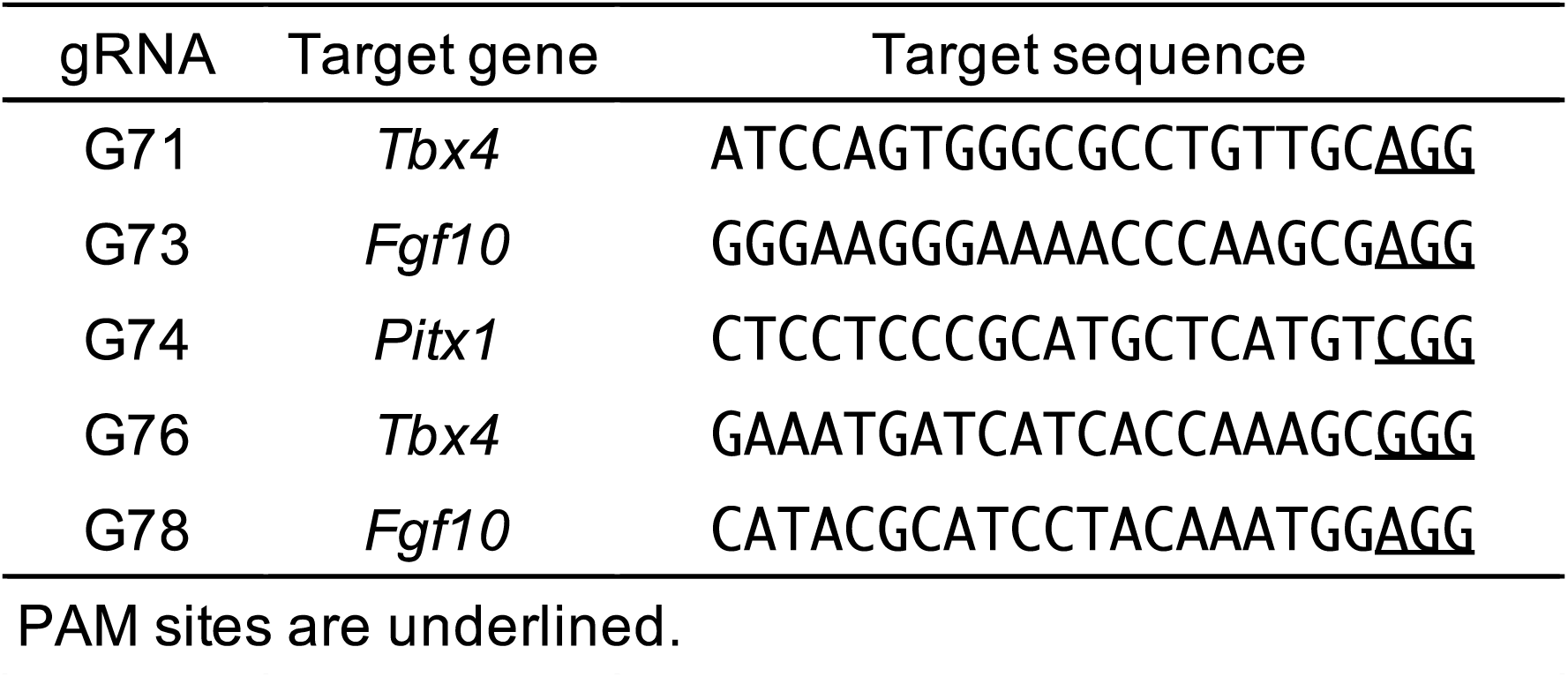
Target sequence of gRNA.

**Table S2.**
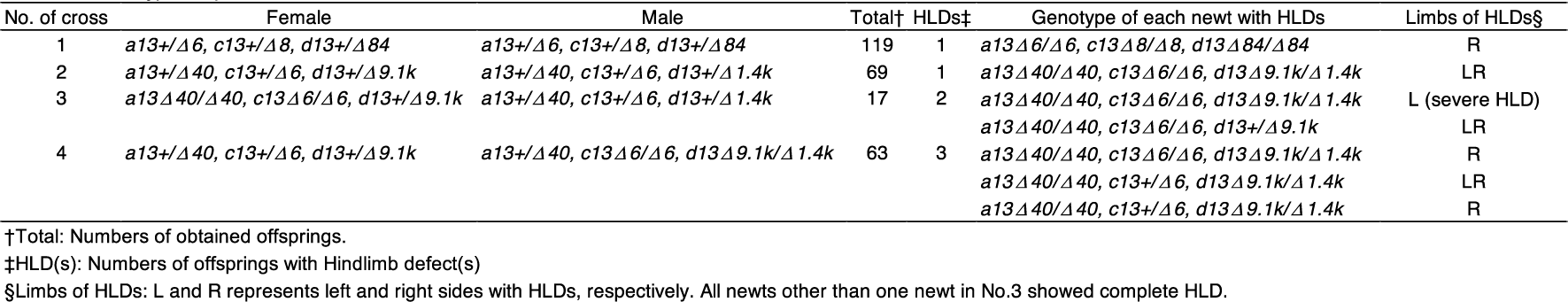
Genotypes of parents and mutants with hindimb defects.

**Table S3.**
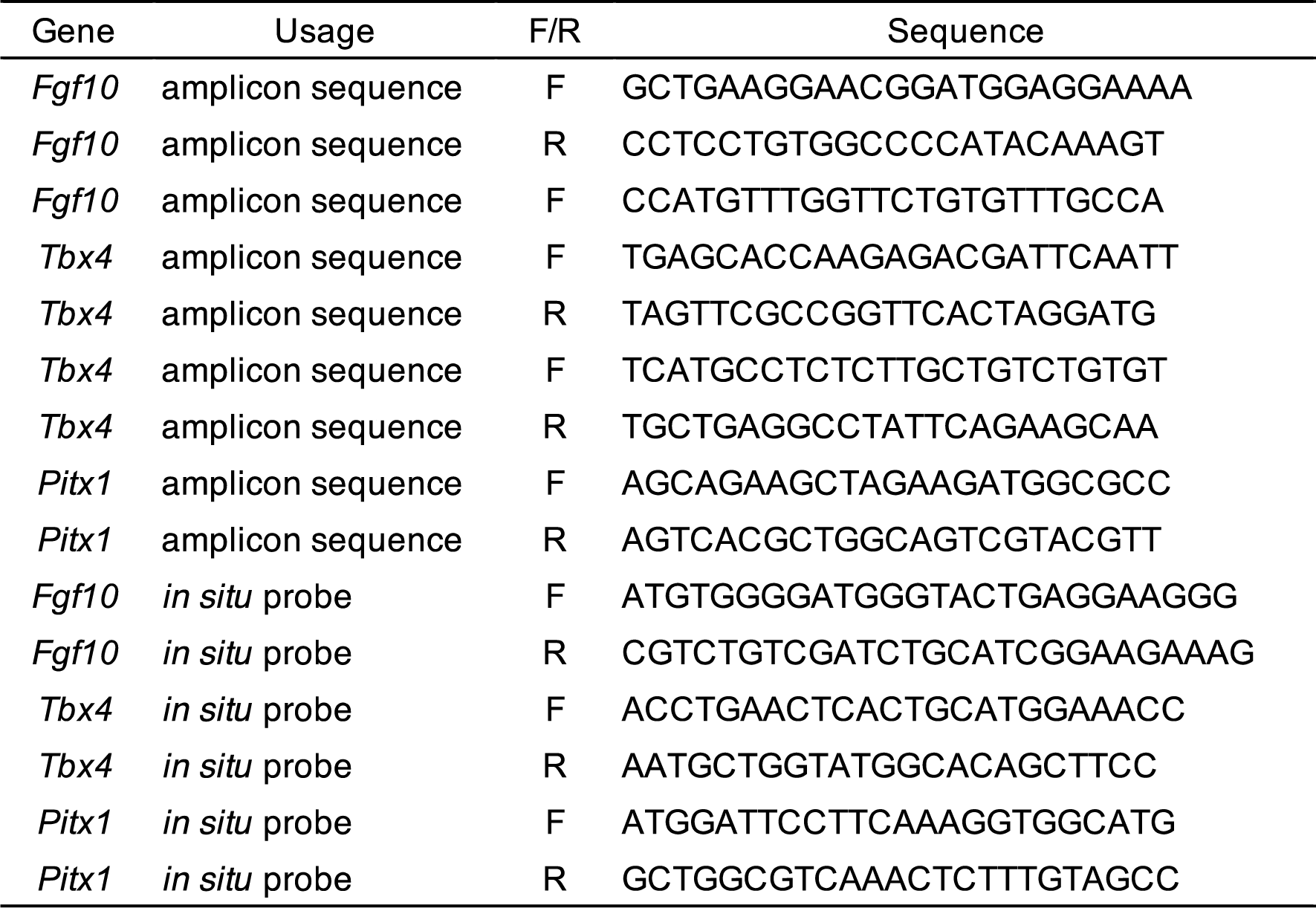
DNA sequence of PCR primers.

**Table S4.**
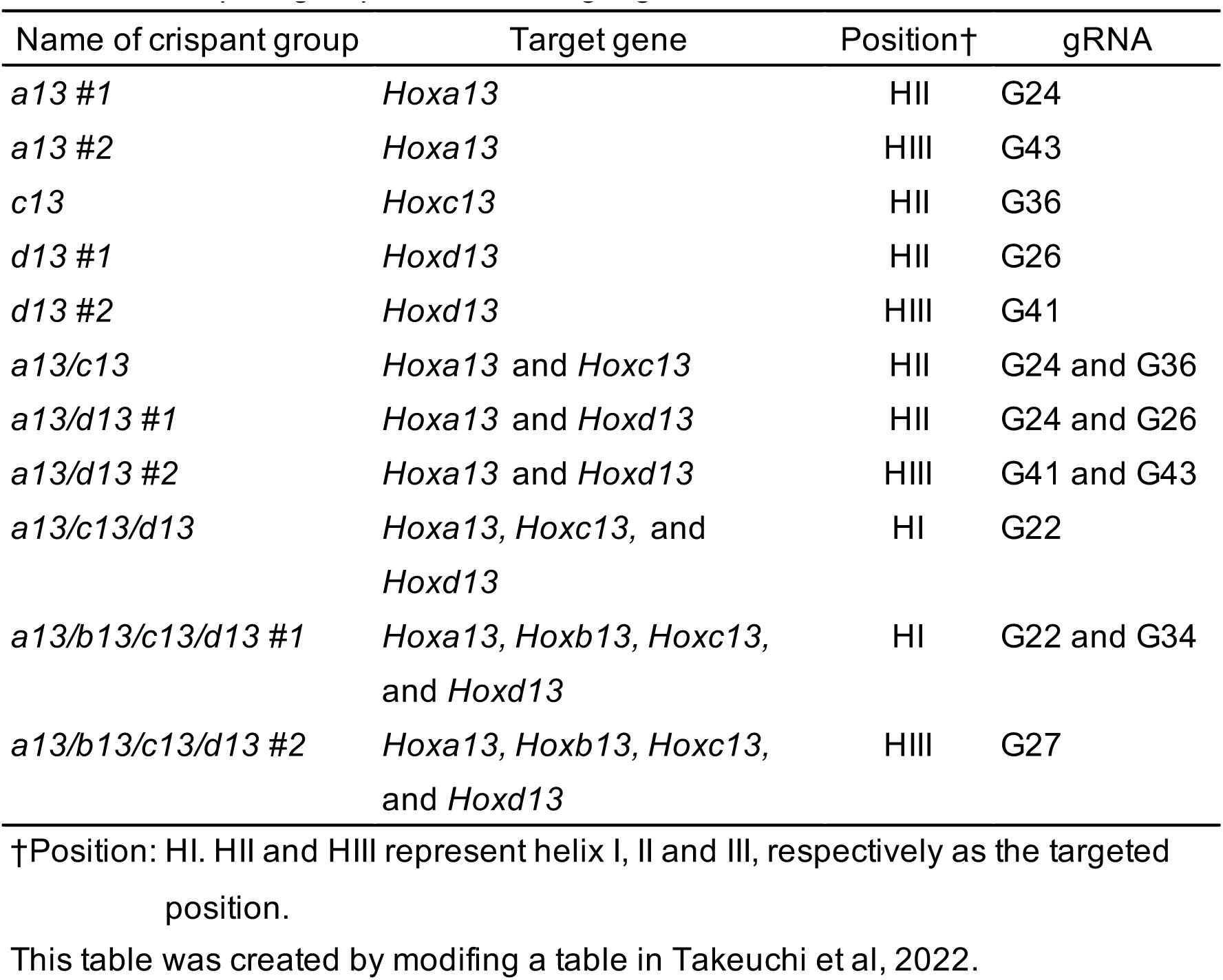
Crispant groups and their target gene I.

**Table S5.** The frequency of hindlimb defects variations.

| Crispant group | Total newt No. | No. of newts with HLDs† |  |  |
| --- | --- | --- | --- | --- |
|  |  | Bilateral | Unilateral (L, R) | Total |
| *a13/d13 #1* | 84 | 13 | 21 (12, 9) | 34 |
| *a13/d13 #2* | 50 | 7 | 10 ( 4, 6) | 17 |
| *a13/c13/d13* | 141 | 33 | 54 (24, 30) | 87 |
| *a13/b13/c13/d13 #1* | 18 | 5 | 2 ( 0, 2) | 7 |
| *a13/b13/c13/d13 #2* | 25 | 5 | 7 ( 3, 4) | 11 |
| Total | 318 | 63 | 93 (43, 51)<br>n.s.‡ | 156 |

†HLDs: hindlimb defects
‡n.s.: not significant in Fischer's exact test.

**Table S6.**
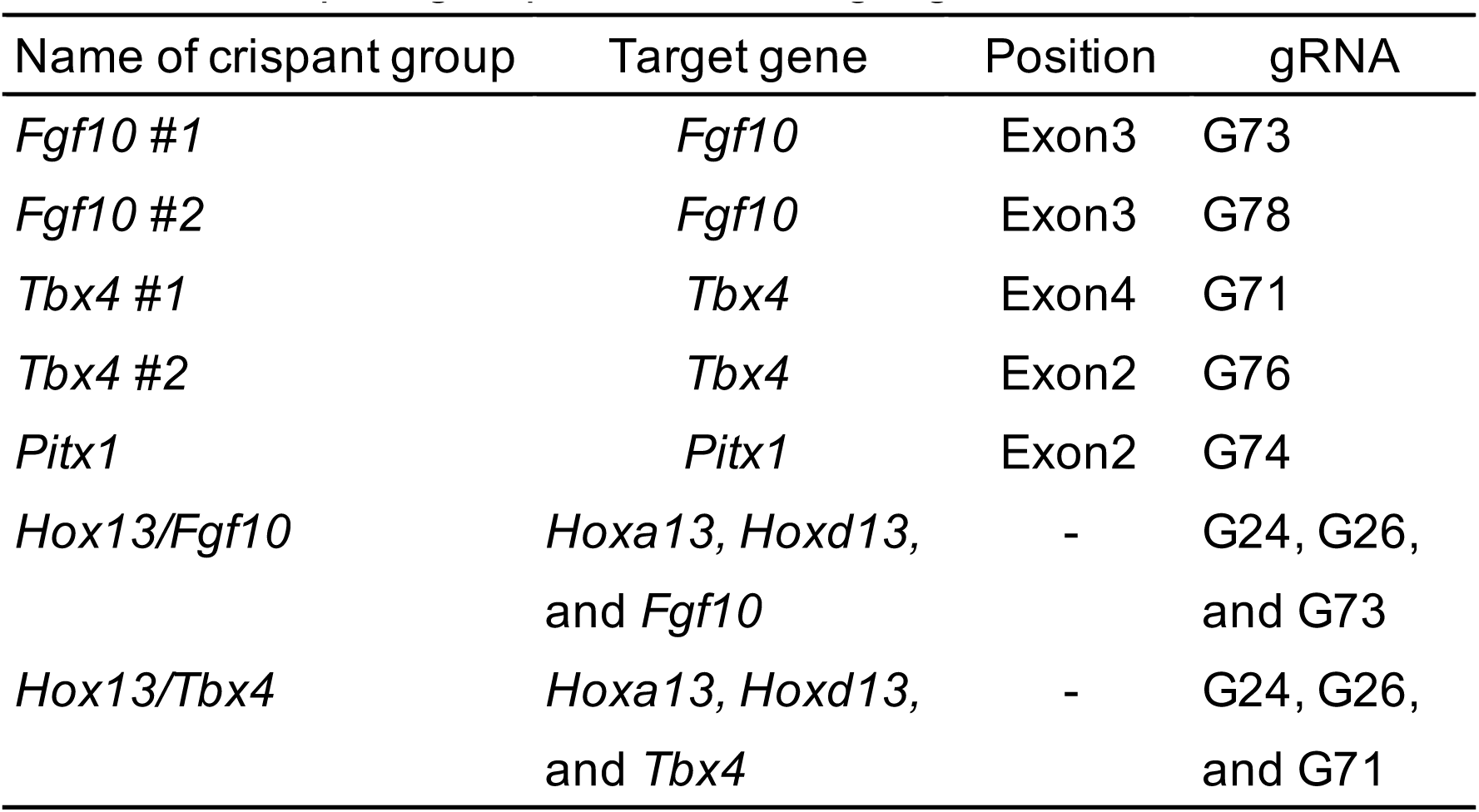
Crispant groups and their target gene II.

| Name of crispant group | Target gene | Position | gRNA |
| --- | --- | --- | --- |
| *Fgf10* #1 | *Fgf10* | Exon3 | G73 |
| *Fgf10* #2 | *Fgf10* | Exon3 | G78 |
| *Tbx4* #1 | *Tbx4* | Exon4 | G71 |
| *Tbx4* #2 | *Tbx4* | Exon2 | G76 |
| *Pitx1* | *Pitx1* | Exon2 | G74 |
| *Hox13/Fgf10* | *Hoxa13*, *Hoxd13*,<br>and *Fgf10* | - | G24, G26,<br>and G73 |
| *Hox13/Tbx4* | *Hoxa13*, *Hoxd13*,<br>and *Tbx4* | - | G24, G26,<br>and G71 |

**Table S7.** Mutation rate of each gRNA.

| Name of crispant group | Analyzed crispants | Mean of mutation rate±SD | Total reads | Avarage reads |
| --- | --- | --- | --- | --- |
| *Fgf10* #1 | 7 | 97.2±5.2 | 32492 | 5415.3 |
| *Fgf10* #2 | 2 | 98.5±1.5 | 89944 | 44972.0 |
| *Tbx4* #1 | 8 | 86.5±8.5 | 124634 | 15579.3 |
| *Tbx4* #2 | 3 | 96.4±2.5 | 95526 | 31842.0 |
| *Pitx1* † | 4 | 90.6±5.1 | 10216 | 2554.0 |
| *Pitx1* ‡ | 3 | 40.7±8.8 | 9585 | 3195.0 |

†: Crispants with abnormal phenotypes
‡: Crispants without abnormal phenotypes

**Table S8.**
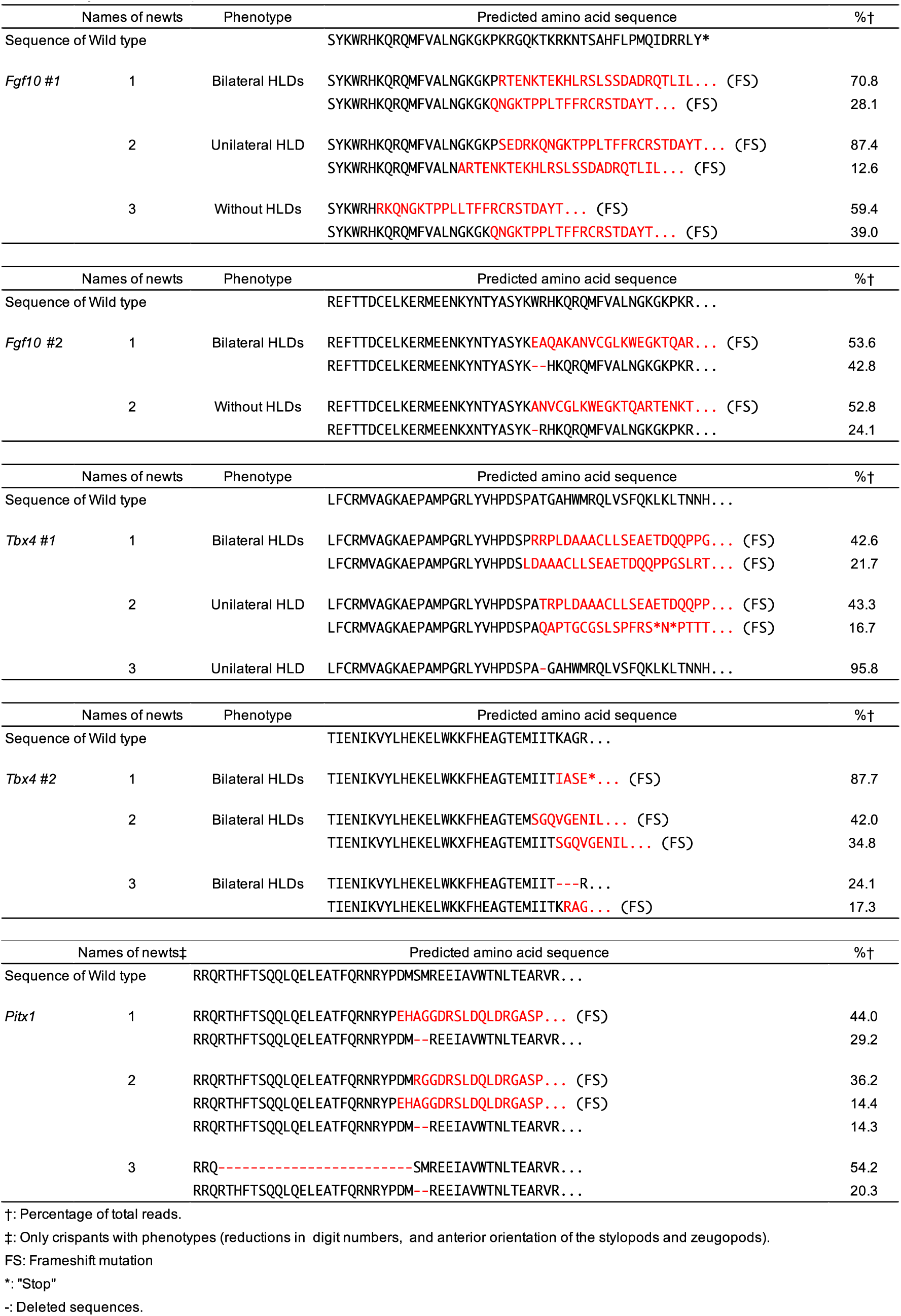
Major alleles of crispants.

**Table S9.** Hindlimb phenotypes of *Fgf10* crispant newts.

| Phenotype | No. of limbs | % |
| --- | --- | --- |
| Normal | 3 | 3.8 |
| 0-4d† | 70 | 89.7 |
| Severe HLD | 2 | 2.6 |
| Complete HLD | 3 | 3.8 |
| Total | 78 | 100 |

*Fgf10* gRNA #2
| Phenotype | No. of limbs | % |
| --- | --- | --- |
| Normal | 40 | 29.9 |
| 0-4d† | 91 | 67.9 |
| Complete HLD | 3 | 2.2 |
| Total | 134 | 100 |

†: Reduction of digit No.

**Figure S1.**
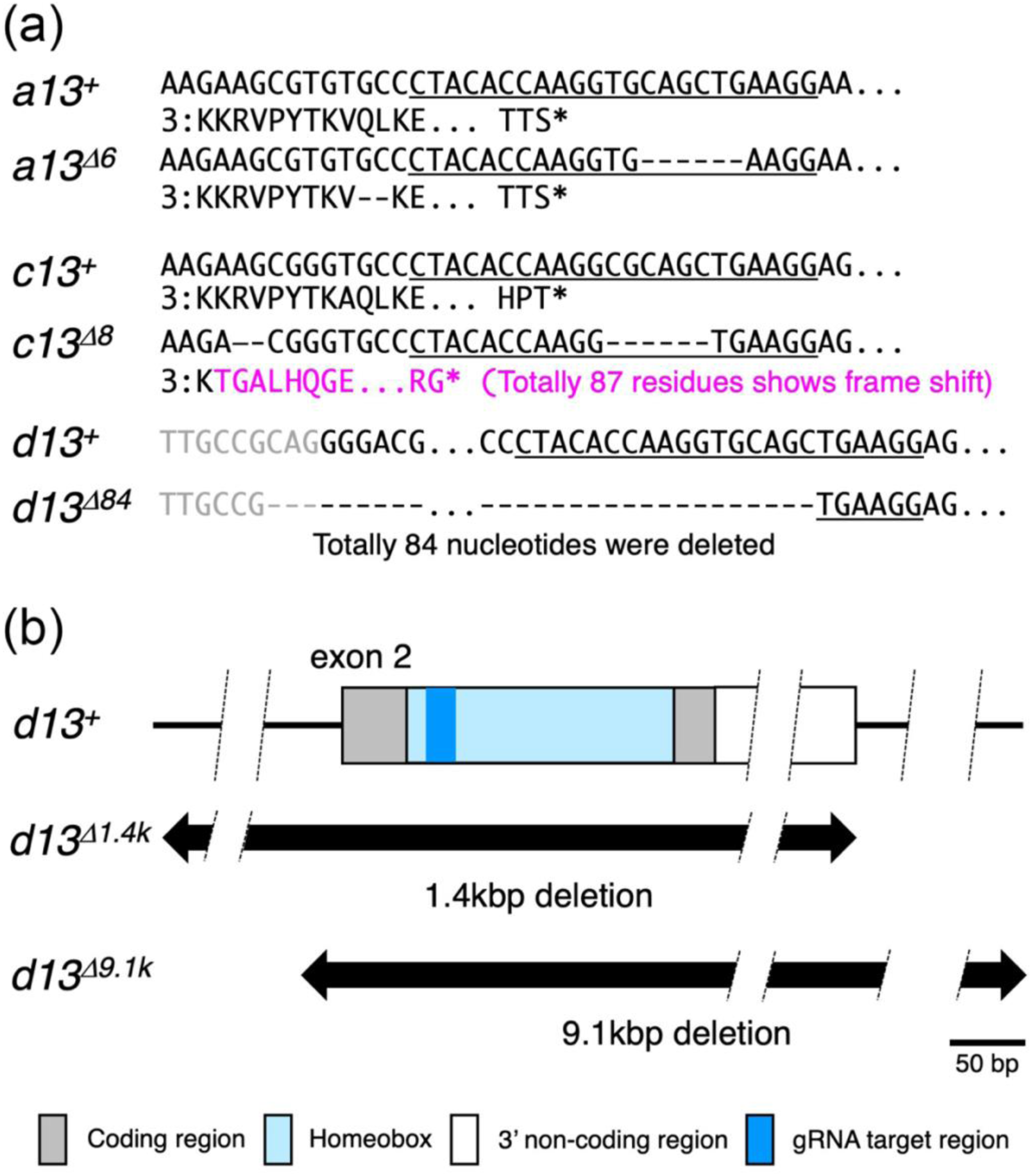
Nucleotide sequences and predicted deletion regions of *Hox13* mutant alleles. (a) Nucleotide and amino acid sequences of *Hoxa1^Δ6^*, *Hoxc13 ^Δ^ ^8^* and *Hoxd13 ^Δ^ ^84^*. Target sequences of gRNA are underlined. The numbers to the left of the amino acid sequences indicate the position of the first residue in the homeodomain. Magenta characters, gray characters, asterisks and dashes indicate sequences different from the wild-type owing to a frameshift, a sequence in the first intron, “stop”, and deleted sequences, respectively. (b) Predicted deletion regions of *Hoxd13 ^Δ^ ^1.4k^* and *Hoxd13 ^Δ^ ^9.1k^*. The exact 3’ end position of the 3’ non-coding region is unknown.

**Figure S2.**
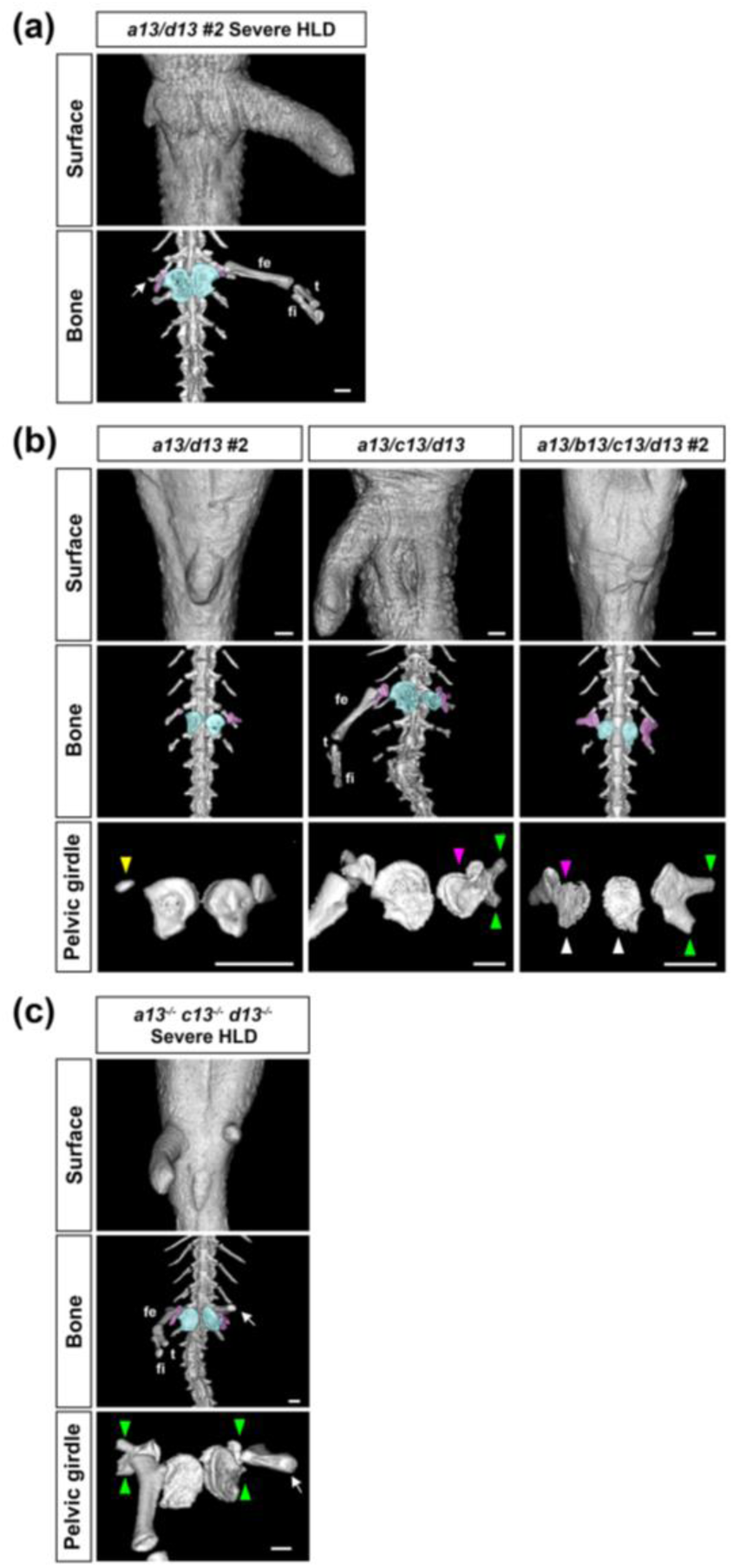
Variations of phenotypes in *Hox13* crispants. Arrows indicate a rudimentary femurs. (a) Micro-CT ventral images of severe HLDs in an *a13/d13 #2* crispant at approximately 7 mpf. (b) Variations in pelvic girdle malformations. Micro-CT ventral images of hindlimbs and pelvic girdles of *Hox13* crispants with HLDs at around 5 mpf. Regions shown by colors and abbreviations are the same as in Figure 1. The yellow, pink, green, and white arrowheads show hypoplasia in ilium, fusion of ischiopubis and ilium, bifurcation in the ilium, and hypoplasia in ischiopubis, respectively. (c) Micro-CT ventral images of hindlimbs and pelvic girdles of *a13^-/-^ c13^-/-^ d13^-/-^* germline mutant with severe HLDs at 6 mpf. The middle panel represents whole bones around the hindlimbs. Regions shown by colors and abbreviations are the same as in Figure 1. Green arrowheads show bifurcation in the ilium. Scale bar, 1 mm.

**FIGURE S3.**
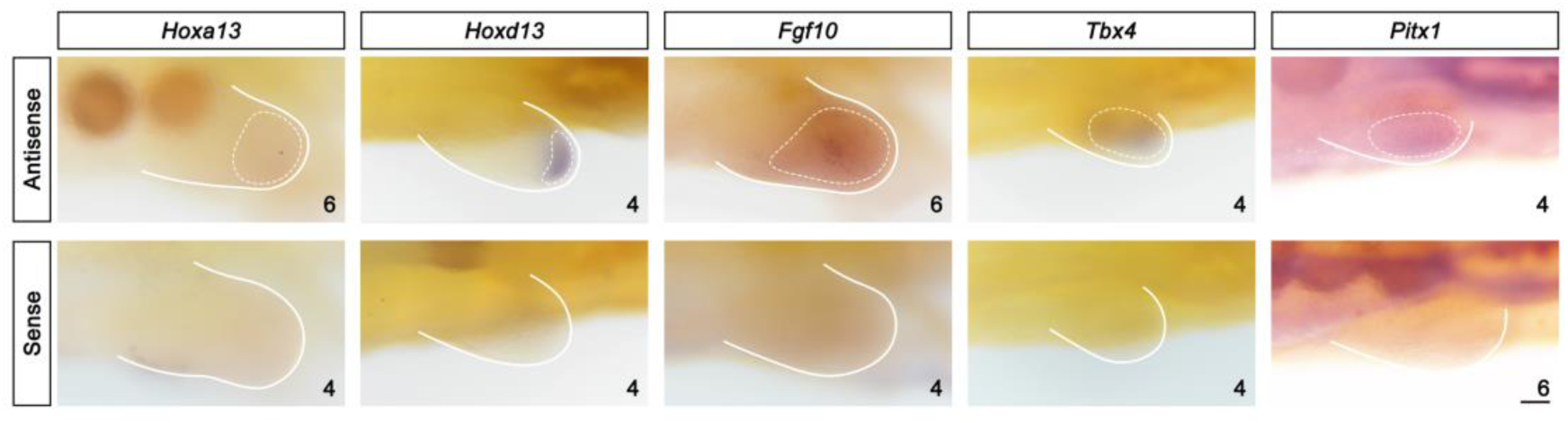
Antisense probe-specific WISH signals in hindlimb buds of wild-type newts. Upper and lower panels show the WISH images of hindlimb buds hybridized with antisense and sense probes, respectively, in wild-type newts. The areas surrounded by the dotted lines indicate antisense probe-specific signals. *Hoxa13*: lateral view, 22dpf; *Hoxd13*: ventral view, 23dpf; *Fgf10*: lateral view, 22dpf; *Tbx4*: ventral view, 24dpf; *Pitx1*: ventral view, 17dpf. The number of hindlimb buds showing similar results is shown in each image. Scale bar, 0.1 mm.

**Figure S4.**
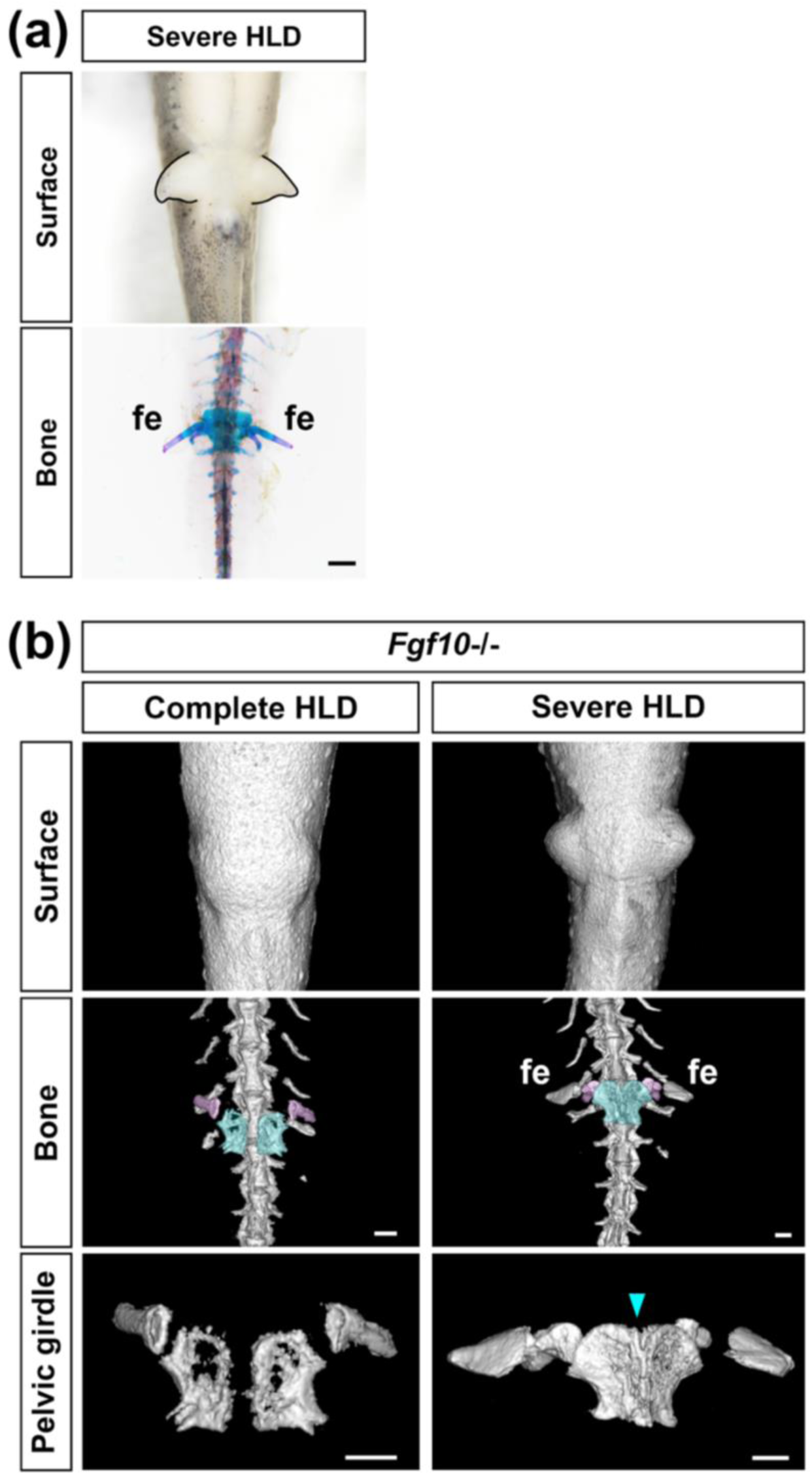
Phenotypes of *Fgf10* mutant newts. (a) Severe HLDs of *Fgf10* #1 crispants at 4 mpf. (b) *Fgf10*-/- germline mutant newts at 10 mpf with complete HLDs or severe HLDs. Light blue arrowheads indicate a fusion of left and right ischiopubis. Regions are shown with the same colors as in Figure 1. Scale bar, 1 mm.

**Figure S5.**
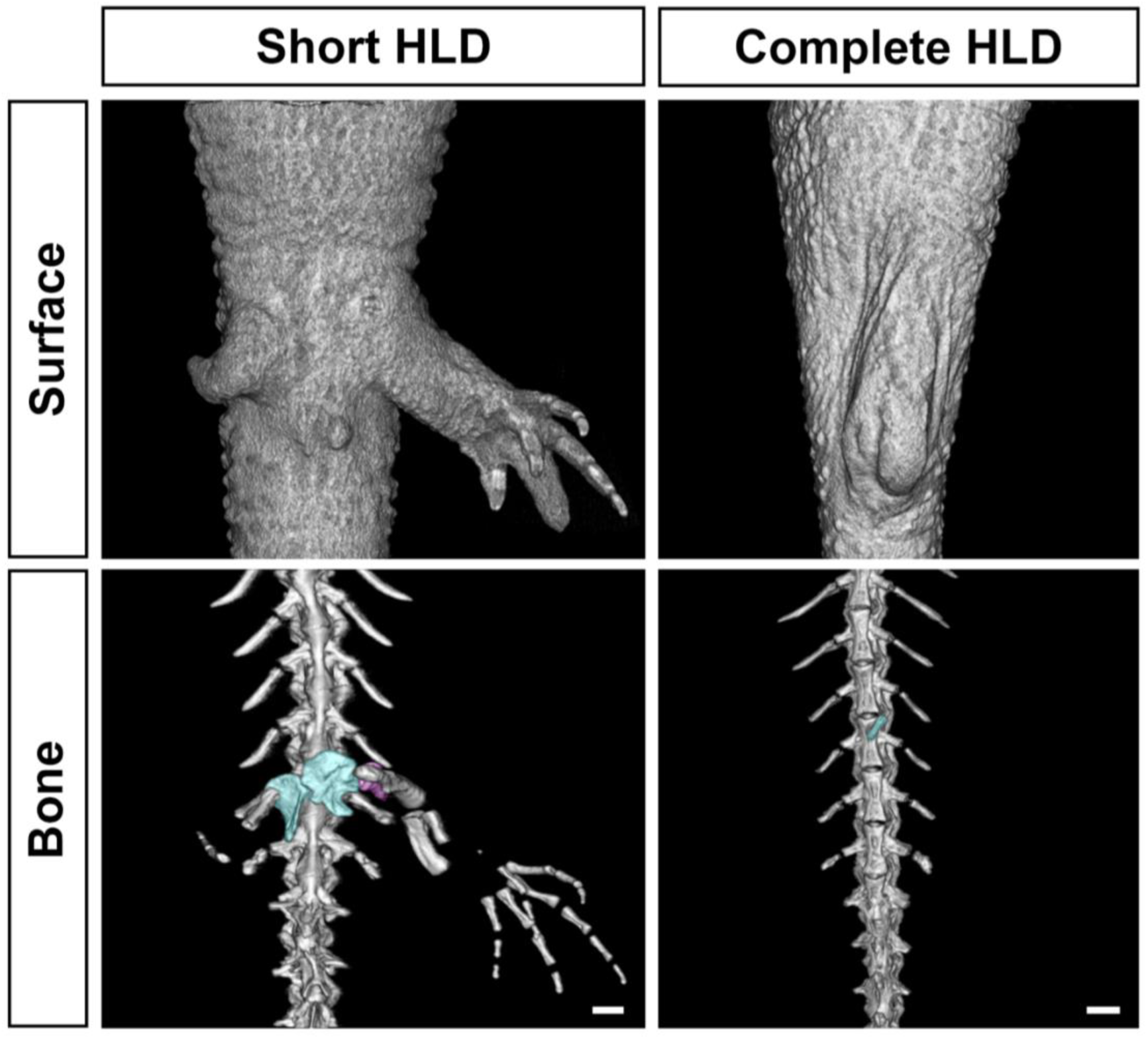
Phenotypic variation in LB+ sides of *Tbx4* #1 crispants. The regions shown in colors are the same as those in Figure 1. Scale bar, 1 mm.

## Notes

### Competing Interest Statement

The authors have declared no competing interest.

